# Sex-specific modifications of gametogenesis in natural and lab-bred *Fundulus* spp. hybrids

**DOI:** 10.1101/2025.11.26.690728

**Authors:** Dmitrij Dedukh, Ksenia Obidena, Zuzana Majtanova, Pascale Ouellette, Alexandre-James Roussel, Marianna E. Horn, Karel Janko, Anne C. Dalziel, Anne-Marie Dion-Côté

## Abstract

Successful interspecific hybridization can be limited by a variety of reproductive barriers, including the emergence of asexual lineages which, while relatively rare in animals, is an example of a strong post zygotic barrier. In vertebrates, asexual reproduction typically arises through gametogenic alterations such as premeiotic genome endoreplication or meiotic failure, yet the relative frequency of and conditions leading to these alterations remain poorly understood. Here, we investigate natural, all female clonal hybrid lineages and laboratory-bred hybrids of both sexes between two North American killifishes, the Banded Killifish (*Fundulus diaphanus*) and Common Killifish (*F. heteroclitus*), to determine the mechanisms underlying and maintaining the emergence of asexual reproduction. By combining cytogenetic, molecular, and gametogenetic analyses, we show that all natural and some lab-bred F1 hybrid females can produce diploid and tetraploid oocytes, with natural hybrids producing a higher proportion of tetraploid eggs. In diploid oocytes, we observed both bivalents and univalents. In contrast, tetraploid oocytes exclusively formed bivalents with normal crossover formation, consistent with premeiotic genome endoreplication restoring proper meiotic pairing and facilitating clonal reproduction. Wild and lab-bred hybrid males do not undergo premeiotic genome duplication and exhibit extensive chromosomal mispairing, leading to aberrant meiotic divisions, elevated apoptosis, and decreased fertility or sterility. Together, these findings highlight sex-specific differences in gametogenesis, paralleling patterns observed in other systems, and emphasize the relative ease with which asexual reproduction can arise in hybrid females, in contrast to the sterility commonly observed in hybrid males.

## Introduction

Hybridization between closely related lineages can produce fertile F1 offspring, with further reproduction leading to variable outcomes including genetic introgression to variable extent, the formation of a hybrid swarm and homoploid hybrid speciation (Gyllensten et al. 1985; Rhymer & Simberloff 1996; Comeault & Matute 2018; Mallet 2008; Stöck et al. 2021; Bhat et al. 2014; Abbott et al. 2013). As genetic divergence between parental lineages increases, hybrids may be more likely to display transgressive phenotypes or hybrid vigour, but when parental lineages diverge past a certain point, hybrid fertility may decline due to disruptions in chromosome pairing during meiosis (Forsdyke 2003; Stelkens & Seehausen 2009; Marta et al. 2023). Indeed, reductions in hybrid fitness and the development of postzygotic reproductive barriers have been shown to correlate with increased genetic divergence (Coyne & Orr 2004; Marta et al. 2023). In species with dimorphic sex chromosomes, genetic incompatibilities disproportionately affect the heterogametic sex, consistent with Haldane’s rule (Haldane 1922; Stöck et al. 2021). Finally, the accumulation of incompatible alleles or chromosomal differences may result in complete hybrid sterility or inviability, completing speciation (Wu 2001; Coyne & Allen Orr 2004; Marta et al. 2023).

In some cases, although speciation is complete, hybridization between sufficiently related species can occur but disrupts sexual reproduction, giving rise to diverse asexual reproductive modes in metazoans (Abbott et al. 2013; Dawley & Bogart 1989; Schön et al. 2009; Stöck et al. 2021). Such asexual hybrids most commonly arise when genetic and chromosomal divergence between parental species is high, but not divergent enough to lead to complete hybrid sterility or inviability (Shaw & Mullen 2014; Stankowski & Ravinet 2021; Stöck et al. 2021). In hybrid vertebrates, alterations in gamete production, which can be either premeiotic or meiotic, can overcome potential hybrid sterility and produce hemiclonal or clonal gametes (Dawley & Bogart 1989; Schön et al. 2009; Neaves & Baumann 2011; Stöck et al. 2021). Further alterations of fertilization mechanisms can lead to the establishment of clonal lineages that reproduce without introgression from either parental genome (Lamatsch & Stöck 2009; Zhang et al. 2015; Dedukh & Krasikova 2022). Such modifications in gametogenesis and fertilization have led to diverse forms of asexual reproduction, which have independently emerged across taxa, such as parthenogenesis (in some fish and reptiles), gynogenesis (in certain fish), and hybridogenesis (in fish and amphibians) (Dawley & Bogart 1989; Schön et al. 2009; Stöck et al. 2021).

In asexual organisms, clonal gametes are produced by a wide range of cytological mechanisms, from entirely ameiotic (apomictic) processes to modified meiosis (automixis) (Stenberg & Saura 2009, 2013; Neaves & Baumann 2011). In invertebrates and in certain cases of facultative parthenogenesis in vertebrates, suppression of first or second polar body extrusion or gamete fusion may contribute to unreduced gamete formation (Stenberg & Saura 2009; Lampert 2008; Blanc et al. 2023). By contrast, only two mechanisms have been observed in asexual hybrid vertebrates: premeiotic genome endoreplication and meiotic failure (ameiosis) (Stenberg & Saura 2009; Stöck et al. 2021). In endoreplication, gonocytes duplicate their genomes (i.e., diploids become tetraploid), enabling proper pairing of replicated chromosomes from the same parental species during meiosis (Cimino 1972; Kuroda et al. 2018; Lutes et al. 2010; Macgregor & Uzzell 1964; Stöck et al. 2002; Dedukh et al. 2020; Dedukh, Altmanová, et al. 2022; Dedukh et al. 2024). In contrast, during achiasmatic meiosis observed in some fish hybrids (i.e., *Poecilia formosa* and *Carassius* hybrids), gonocyte genomes remain unmodified, but they proceed through meiosis without recombination or bivalent formation (Monaco et al. 1984; Dedukh, da Cruz, et al. 2022; Lu et al. 2022). In such cases, the reductional division is skipped, while the equational division proceeds normally, resulting in unreduced eggs (Yamashita et al. 1993). The reasons why certain organisms employ one gametogenic modification, such as meiotic failure or premeiotic endoreplication, rather than another remain unknown. Moreover, these mechanisms appear relatively flexible and may vary depending on ploidy level within a species (Alves et al. 1998; Lafond et al. 2019; Arai 2023; Dedukh et al. 2024; Vukić et al. 2025). Thus, characterizing additional lineages should help disentangle the factors that might result in meiotic failure versus premeiotic genome endoreplication and better understand the range of mechanisms underlying these processes.

In fish hybrids, clonal reproduction has arisen across diverse lineages (reviewed by Dalziel et al. (2020)). One clonal fish for which gametogenesis mechanisms are not yet known is the asexual killifish produced from hybridization between the Common (*Fundulus heteroclitus*) and Banded Killifish (*F. diaphanus)* (Dawley 1992; Dalziel et al. 2020; Hernandez Chavez & Turgeon 2007; Fritz & Garside 1974). These non-sister species are estimated to have diverged ∼ 15-25 MYA, but currently hybridize in brackish waters along the Atlantic Canadian Coast in multiple sites where the freshwater-preferring Banded Killifish and salt-water preferring Common Killifish habitats overlap (Hernandez Chavez & Turgeon 2007; Dawley 1992; Fritz & Garside 1974; Mérette et al. 2009; Ghedotti & Davis 2017). In at least two sites in Nova Scotia, Canada (Porters Lake, Saint Mary’s River Estuary), hybridization leads to the production of asexual, clonal, all-female lineages (Dawley 1992; Hernandez Chavez & Turgeon 2007; Dawley et al. 1999). In this paper, we first aim to characterize the type of gametogenic alteration occurring in natural hybrid lineages (premeiotic endoduplication vs. meiotic failure).

Gametogenic alterations causing clonal gametes formation are generally thought to be rare and restricted to a single event, as in *P. formosa* (Stöck et al. 2010; Warren et al. 2018). Nevertheless, emerging data from other vertebrate asexual complexes has shown that gametogenic alterations leading to asexuality, such as premeiotic genome endoreplication, can emerge spontaneously and relatively frequently in F1 offspring of inter-specific crosses (Marta et al. 2023; Dedukh et al. 2021; Dedukh et al. 2024; Choleva et al. 2023). Moreover, premeiotic genome endoreplication can be induced in artificial F1 hybrids from species that do not naturally hybridize in the wild (*Cobitis*, *Clarias*, Medaka) (Shimizu et al. 2000; Marta et al. 2023; Dedukh et al. 2024). Indeed, in *Fundulus*, multiple clonal lineages have been detected in the wild and there is evidence new lineages may continue to form (Hernandez-Chavez and Turgeon, 2007; Dalziel et al. 2020). As well, lab-produced F1 hybrids from both female *F. diaphanus* × male *F. heteroclitus* and female *F. heteroclitus* × male *F. diaphanus* crosses can develop to adulthood (Fritz and Garside 1974a). Moreover, female *F. diaphanus* × male *F. heteroclitus* F1 lab-bred hybrid females also produce eggs (pers. obs.); however, it remains uncertain whether these eggs are clonal or not. Our second aim was to test whether gametogenic alterations leading to asexual reproduction may emerge *de novo* in lab-bred reciprocal F1 hybrids.

Notably, asexuality tends to emerge in hybrid females, while males are generally sterile or at least unable to reproduce asexually (with a few exceptions, such as in cases of androgenesis, *Pelophylax* water frogs, or *Bufotes* toads) (Graf 1989; Komaru et al. 1998; Stöck et al. 2002; Spangenberg et al. 2017; Kuroda et al. 2019; Dedukh et al. 2020, 2024; Marta et al. 2023). Because wild hybrid males are rarely found in one hybrid zone in Porters Lake, NS (Fritz & Garside 1974; Dalziel et al. 2020; Hernandez Chavez & Turgeon 2007; Mérette et al. 2009), our third aim was to determine whether F1 hybrid males can produce gametes and whether they can reproduce asexually.

## Materials and Methods

### Wild fish collection and lab crosses

All experimental procedures with fish were approved by local Animal Care Committees at Université de Moncton or Saint Mary’s University (A-M.D-C.: protocols 19-02 and 22-02, A.C.D.: protocol 20-06). Wild *F. diaphanus*, *F. heteroclitus* or asexually reproducing hybrids were collected over the summers of 2020, 2022, 2023 and 2024 from Porters Lake, NS (44.68414, −63.30238) under a Canadian Department of Fisheries and Oceans Scientific Fish collection permit to A.C. Dalziel (343930) and kept in stand-alone brackish water tanks (10-12 ppt) at 18-21°C until they were bred or euthanized.

Fish sex and species were initially determined in the field based upon breeding colouration and general morphology as outlined by MacPherson et al. (2023), and putative hybrids were identified using the set of three morphological measurements described by Mérette (2009). All fish used in breeding and chromosomal preparations were genotyped (see “**DNA extraction and genotyping**”) to confirm species identity and determine maternal ancestry. Herein, genotypes are reported in the format *female × male*. For example, D×D refers to pure crosses with a *F. diaphanus* mother and father while D×H indicates F1 hybrids with a *F. diaphanus* mother and *F. heteroclitus* father. The vast majority (>90%) of hybrids at Porters Lake are F1 clonal D×H females (MacPherson et al. 2023; Hernandez Chavez & Turgeon 2007; Mérette et al. 2009; Dawley et al. 1999) and all wild female hybrids (n=6) used in this study had a *F. diaphanus* mother and *F. heteroclitus* father.

We sampled lab-bred F1 hybrids bred in 2020 (MacPherson et al. 2023) and 2023, all of which were euthanized in July-August 2024. The 2020 F1 crosses were approximately four years old at time of sampling and included seven adult F1 hybrid males from mixed brood families (three D×H and four H×D individuals from an unknown number of parents) and one adult D×H F1 hybrid female. The 2023 F1 crosses included pure and reciprocal hybrid juvenile males and females, which were approximately one year old at the time of sampling and were not sexually mature. The 2023 lab-bred F1 fish consisted of offspring from seven pooled D×D crosses, two pooled D×H crosses, six pooled H×D crosses and five pooled H×H crosses. Table 1 summarizes the individuals analyzed by method (i.e., FISH, CGH, pachytene analysis and gonadal microanatomy - see below). As families within a given cross type were pooled, the family of origin of each individual is not known.

**Table 1.**
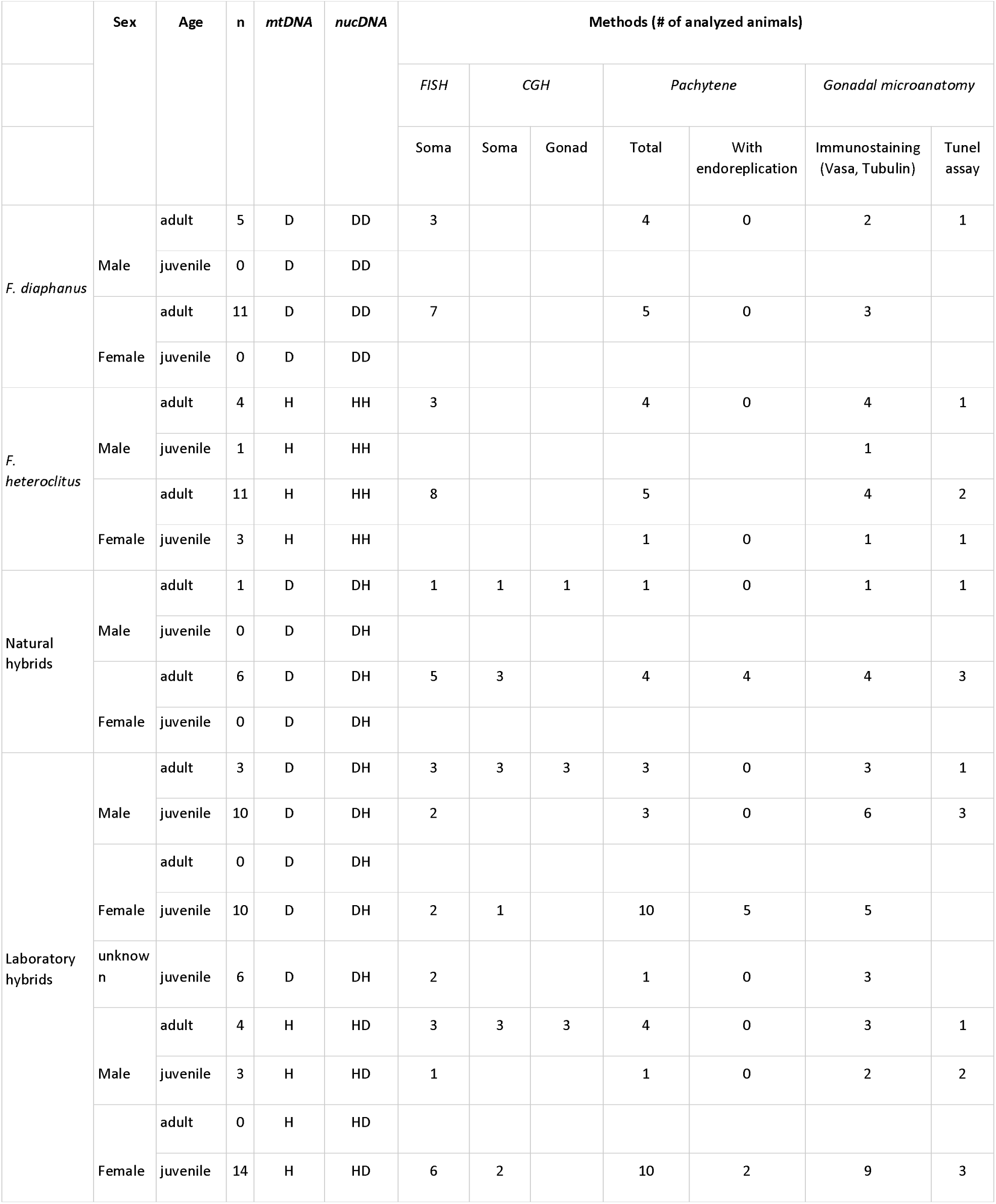
Summary of the animals used in this study and the methods applied. mtDNA stands for mitochondrial genotype determined by RFLP whereas nucDNA means nuclear genotype as determined by microsatellite analysis. All juveniles were lab-bred in 2023.

### DNA extraction and genotyping

DNA was extracted from liver or fin clips using the E.Z.N.A tissue DNA extraction kit (Omega Bio-tek). We amplified a D-loop mtDNA fragment to determine maternal ancestry using a restriction length fragment polymorphism (RFLP) assay and genotyped two microsatellite loci (FhCA-1 and FhCA-21) to assess species or hybrid status as previously described (Dalziel et al. 2020). Microsatellite allelic sizes were determined by capillary electrophoresis on a 3730xl DNA Analyzer at The Center for Applied Genomics in Toronto and analyzed using the microsatellite application (MSA) from the Thermo Fisher Cloud.

### Summary of individuals analyzed and techniques used

In total, we investigated the following groups of wild fish: 16 pure *F. diaphanus* individuals (11 females, 5 males) and 19 pure *F. heteroclitus* individuals (14 females, 5 males), along with 7 natural asexual hybrids (6 D×H females, 1 D×H male). Additionally, we analyzed lab-bred F1 hybrids: 29 D×H individuals (10 females, 13 males, 6 juveniles with unknown sex from two pooled families) and 21 H×D individuals (14 females, 7 males from six pooled families) (Table 1).

We examined mitotic and meiotic chromosomes for all individuals, including synaptonemal complexes during pachytene (males and females) and meiotic metaphase chromosomes (males only due to technical limitations). Mitotic metaphase chromosomes were further analyzed for hybrid individuals by comparative genomic hybridization (CGH) and fluorescence *in situ* hybridization (FISH) with newly developed satellite markers (see “**Identification of species-specific satellite repeats and fluorescence *in situ* hybridization (FISH)**”). We analyzed gonadal microanatomy using confocal microscopy in sexually reproducing males and females and both lab-bred and wild hybrids.

### Mitotic and meiotic metaphase chromosome preparation

Mitotic metaphase chromosome spreads were prepared from gills and kidneys of adult and juvenile pure *F. diaphanus* (7 females, 3 males), pure *F. heteroclitus* (8 females, 3 males), natural hybrids (5 females, 1 male), and F1 lab-bred D×H hybrids (2 females, 5 males, 2 individuals with unknown sex) and F1 lab-bred H×D hybrids (6 females and 4 males). Meiotic metaphase chromosomes were prepared from testes of pure-bred, sexual *F. diaphanus* (3 males), pure-bred, sexual *F. heteroclitus* (3 males), a natural hybrid (1 male), F1 lab-bred D×H hybrids (3 males) and F1 lab-bred H×D hybrids (3 males) following standard protocols (Bertollo et al. 2015).

Adults (wild fish and 2020 lab-bred F1 hybrids) were injected with 0.3 ml of 0.04% colchicine solution, while juveniles (2023 lab-bred F1 hybrids) were kept in 0.1% colchicine solution for 3-4 hours. After dissection, the kidneys and testis fragments were immersed in distilled water for 30 minutes, then fixed in a 3:1 ethanol-to-glacial acetic acid solution, which was replaced three times. The fixed tissues were stored at 4°C until use. To obtain the cell suspension, tissue fragments were placed in 70% glacial acetic acid for one minute and thoroughly macerated. The resulting suspension was dropped onto preheated slides at 60 °C and evenly distributed across the surface. Excess liquid was removed, allowing interphase nuclei and metaphase chromosomes to remain on the slide after evaporation. Metaphase chromosomes were stained with a 5% Giemsa solution for 10 minutes at room temperature to assess chromosome number, morphology, and bivalent formation.

### Synaptonemal complex spread preparation and immunofluorescent staining

Synaptonemal complexes (SC) spread during the pachytene stage of meiosis were prepared from adult male testes (wild fish and 2020 lab-bred F1 males) following the protocols of Moens (2006) and from ovaries of juvenile individuals (2023 lab-bred F1 males and females) according to Araya-Jaime et al. (2015). A small fragment of testis was homogenized in 200 µl 1× PBS. Subsequently, 1 µl of the cell suspension was added to a 30 µl drop of hypotonic solution (0.33× PBS) previously placed on SuperFrost® slides. After 20 minutes, cells were fixed in 2% paraformaldehyde for 4 minutes and air-dried. Ovarian tissues were homogenized in 0.2 M sucrose, and the resulting cell suspension (20 µl) was dropped onto SuperFrost® slides (Menzel Gläser), followed by the addition of 40 µl of 0.2 M sucrose and 40 µl of 0.2% Triton X-100 for 7 minutes. Cells were then fixed by adding 400 µl of 2% paraformaldehyde for 16 minutes, and air-dried. Dried slides with testicular and ovarian SC spreads were placed in 1× PBS and used for immunofluorescent staining.

To detect SC components, we used rabbit polyclonal antibodies (ab15093, Abcam) against SYCP3 to detect lateral elements, and chicken polyclonal antibodies against SYCP1 (a gift from Sean M. Burgess (Blokhina et al. 2019)) to detect the transversal element. The combination of α-SYCP3 and α-SYCP1 allows differentiation between bivalents and univalents, as SYCP1 accumulates only on bivalents, whereas SYCP3 accumulates on both (Blokhina et al. 2019; Dedukh et al. 2020). Crossover loci were identified using antibodies against MLH1 (ab14206, Abcam), and DNA double-strand breaks were visualized by antibodies against RAD51 (GTX00721, GeneTex). Fresh slides with SCs were treated by 1% BSA (Roche) in 1× PBS, 0.01% Tween-20 for 20 minutes, followed by incubation with the primary antibody for 2 hours at room temperature. After three washes in 1× PBS, slides were incubated with secondary antibodies Alexa-488-conjugated goat anti-rabbit IgG (HL+LL) (A-11008, Thermo Fisher Scientific), Alexa-594 goat anti-chicken IgY (HL+LL) (A-11042, Thermo Fisher Scientific) and Alexa-594-conjugated goat anti-mouse IgG (HL+LL) (A-11005, Thermo Fisher Scientific) for 1 hour at room temperature. Finally, slides were washed in 1× PBS and mounted with Vectashield/DAPI (1.5 mg/ml) (Vector, Burlingame, Calif., USA).

### Identification of species-specific satellite repeats and fluorescence *in situ* hybridization (FISH)

Species-specific satellite repeats were identified using the RepeatExplorer2 pipeline (Novák et al. 2013) using established procedures as described elsewhere (Roussel et al. 2025). Briefly, 7 female individuals from Porters Lake (3 *F. heteroclitus*, 4 *F. diaphanus*) were sequenced at low coverage using Illumina sequencing. Reads were then subsampled to ∼0.5X genome-wide coverage and clustered by sequence similarity using RepeatExplorer2. Repeats were quantified for all individuals using RepeatMasker (Smit et al. 2015). We then selected the consensus sequence of eight repeat clusters for which the proportion of reads in the cluster differed by at least 10-fold between *F. heteroclitus* and *F. diaphanus* (Supplementary Material S1).

We used two types of probes against selected satDNA sequences. First, ∼50 bp fragments from each repeat were commercially synthesized and directly labeled with biotin-16-dUTP or digoxigenin-11-dUTP (Supplementary Material S1). Second, PCR products were labeled during amplification with biotin-16-dUTP or digoxigenin-11-dUTP (Roche Applied Science). Primers for PCR were designed within the satellite monomers using Primer-BLAST (Supplementary Material S1).

Fluorescent *in situ* hybridization (FISH) was performed according to Marta et al. (2020). Slides with mitotic and meiotic metaphase chromosomes were treated with 0.01% pepsin/0.01 M HCl for 10 minutes, washed in 1× PBS, dehydrated in ethanol rows (50%, 70% and 96%), and air dried. The hybridization mixture included 40% formamide, 10% dextran sulfate, 2× ЅЅС, 5 ng/μl labeled probe, and 10–50-fold excess of salmon sperm DNA. Probes were denatured at 86 °C for six minutes and placed on ice for 10 minutes. Simultaneously, slides with metaphase chromosomes were denatured at 73 °C for three minutes, immediately transferred through ice-cold ethanol row (70%, 80% and 96%) and air-dried. The denatured probe was applied to slides, covered with cover slides, and incubated overnight at room temperature (RT) in a humid chamber. After hybridization, slides were washed three times in 2× Saline Sodium Citrate buffer (SSC) at 44 °C for 5 minutes each. Streptavidin-Alexa 488 (Invitrogen, San Diego, Calif., USA) and anti-digoxigenin-rhodamine (Invitrogen, San Diego, Calif., USA) were applied to detect biotin-dUTP and digoxigenin-dUTP, respectively. Chromosomal DNA was counterstained with Vectashield/DAPI (1.5 mg/ml) (Vector, Burlingame, Calif., USA).

### Comparative genome hybridization

Comparative genome hybridization (CGH) was conducted on mitotic and meiotic metaphase spreads from natural (n=7) and lab-bred (n=10) F1 hybrid individuals following the protocol of Majtánová et al. (2016). Probes for CGH experiments were prepared using whole genomic DNA (gDNA) from the pure parental species *F. diaphanus* and *F. heteroclitus*. Genomic DNA from *F. diaphanus* and *F. heteroclitus* was extracted using the Qiagen DNeasy Blood & Tissue Kit. The *F. diaphanus* genome DNA was labeled with biotin-16-dUTP (Roche), while the *F. heteroclitus* genome DNA was labeled with digoxigenin-11-dUTP (Roche) using a Nick Translation Mix (Abbott) according to the manufacturer’s protocol. Optimal results were obtained after 2.5 hours of nick translation, yielding labeled DNA fragments approximately 200-500 bp in length.

Nick-translated products from both parental species (1µg of labelled probe for each species) were mixed and dissolved in a hybridization mixture containing 50% formamide, 2× SSC, 10% dextran sulfate, and salmon sperm DNA. Denaturation of slides and probes was performed as described in the FISH protocol. Hybridization was carried out at 37°C for 48 hours, followed by three washes in 50% formamide/2× SSC at 42°C for 5 minutes each, and three washes in 2× SSC at 42°C for 5 minutes each. Probe detection was carried out as described in the FISH protocol. Chromosomes were counterstained with Vectashield/DAPI (1.5 mg/ml) (Vector, Burlingame, CA, USA).

### Whole-mount immunofluorescence staining and TUNEL assay

Before staining, gonadal fragments were permeabilized in 0.5% Triton X-100 in 1× PBS for 4–5 hours at room temperature (RT), then washed in 1× PBS. Terminal deoxynucleotidyl transferase dUTP Nick-End Labeling (TUNEL) assay was applied on gonadal tissue fragments to assess apoptosis following manufacturer’s protocol (ab66110, Abcam).

Whole-mount immunofluorescent staining was performed following the protocol of Dedukh et al. (2021). For immunofluorescent staining, samples were incubated in a blocking solution (1% blocking reagent (Roche) in 1× PBS) for 1–2 hours before being transferred to a fresh blocking solution containing primary antibodies. Mouse monoclonal antibodies against tubulin (ab7291; Abcam) and rabbit polyclonal antibodies DDX4 (C1C3, GeneTex Inc.) against Vasa protein were used as primary antibodies. After overnight incubation at room temperature with primary antibodies, tissues were washed in 1× PBS. Secondary Alexa-488-conjugated goat anti-rabbit IgG (HL+LL) (A-11008, Thermo Fisher Scientific) and Alexa-594-conjugated goat anti-mouse IgG (HL+LL) (A-11005, Thermo Fisher Scientific) were applied overnight at RT. After immunofluorescent staining and TUNEL assay, tissues were stained with DAPI (1 µg/µl; Sigma) overnight in 1× PBS at RT.

### Wide-field, fluorescence and confocal laser scanning microscopy

Slides with mitotic and meiotic metaphase chromosomes were examined on a ZEISS Axio Imager.Z2 epifluorescence microscope equipped with Olympus DP30BW CCD camera and a CoolCube 1 camera (MetaSystems). The IKAROS and ISIS imaging programs (Metasystems) were used to analyze grey-scale images. The captured digital images from FISH experiments were pseudocolored (blue for DAPI, red for anti-digoxigenin-rhodamine, green for streptavidin-FITC) and superimposed using Microimage and Adobe Photoshop.

Slides with SC and diplotene chromosome spreads were analysed on a Provis AX70 Olympus microscope equipped with a standard fluorescence filter set. Chromosome microphotographs were acquired with a CCD camera (DP30W Olympus) using the Olympus Acquisition Software.

Confocal laser scanning microscopy was performed using a Leica TCS SP5 microscope, integrated with an inverted Leica DMI 6000 CS microscope (Leica Microsystems). Specimens were analyzed with an HC PL APO 63× objective. Fluorescent signals were excited using diode and argon lasers for DAPI, Alexa-488 and Alexa-594 fluorochromes, respectively. Images were captured and processed using LAS AF software (Leica Microsystems).

### Statistical analyses

Prevalence of endoreplication, of proper alignment in bivalents, and of bivalent misalignment in female hybrids were assessed statistically (Supplementary Table 1). Females were assessed independently due to differences in sex chromosomes and the absence of endoreplication in males. The proportion of pachytene oocytes that exhibited endoreplication, the proportion of cells in which all bivalents were properly aligned, and the average number of properly aligned bivalents per cell were calculated for each individual. A Shapiro-Wilks test was used to assess normality. Non-normally distributed data (the proportion of oocytes that exhibited endoreplication) were analysed using a Kruskal-Wallis non-parametric test followed by pairwise Mann-Whitney U tests. For normally distributed data (proportion of properly aligned bivalents; average number of alignments per cell), one-way ANOVA was used to compare hybrid genotypes, and with Bonferroni post hoc test to accommodate variable sample sizes (n=4-10). All analyses were performed using SPSS Statistical Software [Version 17.0] (IBM).

## Results

### Wild and lab-bred hybrids have intermediate karyotypes with minor rearrangements in clonal hybrids

We found that the somatic cells (gills, kidneys) of both *F. diaphanus* and *F. heteroclitus* have a diploid chromosome number of 2n = 48, consistent with previous findings (Figure 1) (Dalziel et al. 2020; Arcement & Rachlin 1976; Chen 1970, 1971). We also observed 2n = 48 chromosomes in somatic cells of lab-bred F1 hybrids (D×H and H×D, males and females) and wild asexual hybrids (D×H females) (Figure 1, 2). Both parental chromosome sets were clearly distinguishable in all examined hybrid somatic cells using Comparative Genomic Hybridization (CGH) (Figure 2A, B). Intense signals were observed in centromeric and subtelomeric regions of both parental chromosome sets, suggesting differences in repetitive DNA composition of these regions. Some chromosomes also displayed strong signals in interstitial regions, indicating the presence of species-specific sequences that may correspond to tandem repeats. We did not observe any rearranged or recombined chromosomes in wild asexual female hybrids, indicating that the parental genomes remain structurally intact during clonal propagation.

**Figure 1.**
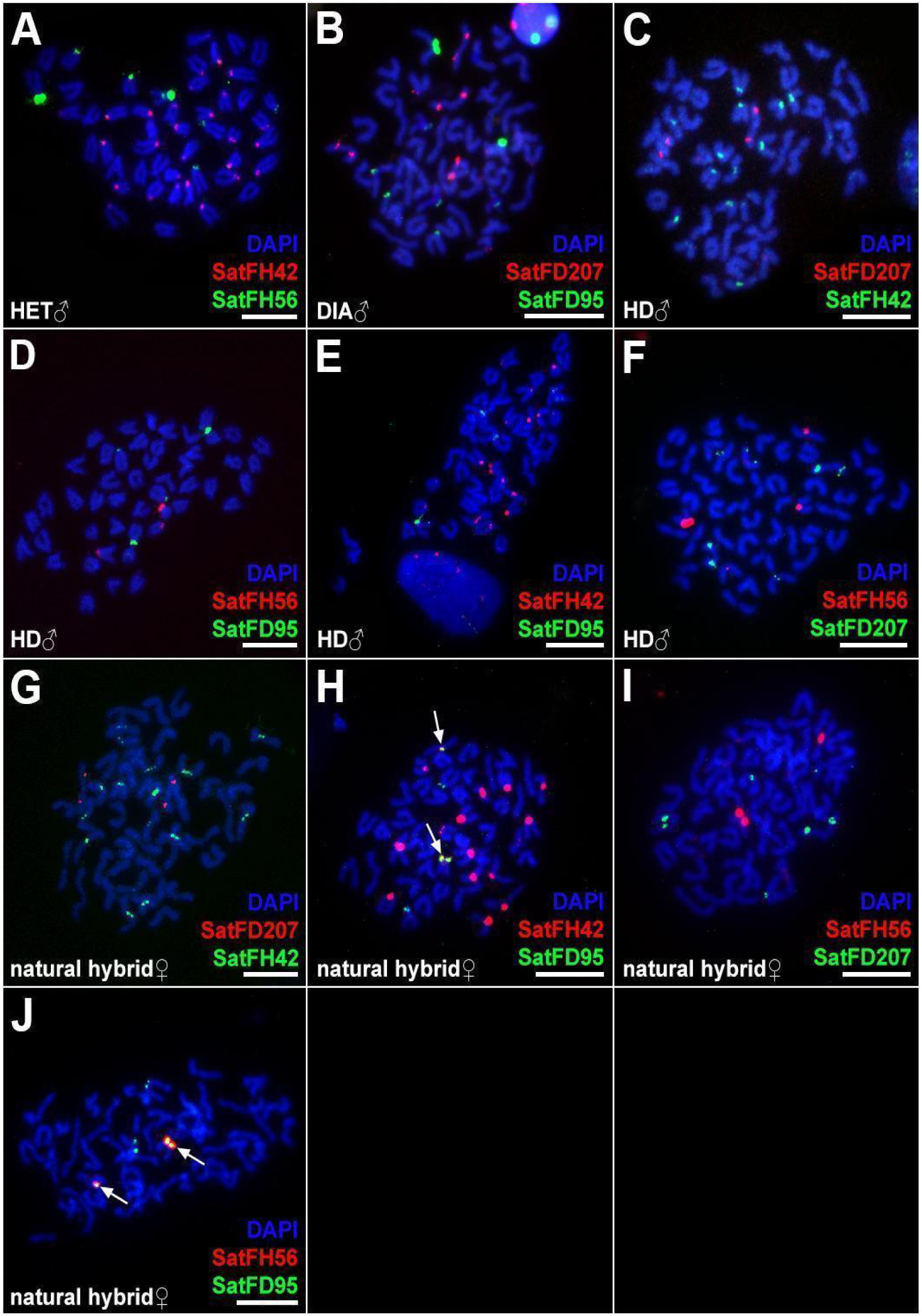
Species-specific satellite DNA distribution in parental *Fundulus* species (A, B), F1 hybrids (C-F), and natural asexual hybrids (G-J) revealed by FISH. Double-colored FISH shows the position of SatFH42, SatFH56, satFD207 and SatFD95 on mitotic chromosomes of corresponding sexual species and their hybrids. (A) In *F. heteroclitus*, SatFH42 shows 12 strong and four weak signals in the pericentromeric regions of 16 chromosome pairs; SatFH56 shows 24 bright signals and eight weak signals in the centromeric regions of 16 chromosome pairs. (B) In *F. diaphanus,* SatFD95 shows four strong and six weak signals in the pericentromeric regions of five chromosome pairs; SatFD207 shows bright signals in the interstitial regions on two chromosome pairs, with weak signals in the subtelomeric region of two chromosome pairs. Diploid F1 HD hybrids (C-F) display exactly half the number of each SatDNA marker, consistent with the haploid genome composition inherited from each parental species. In asexually reproducing wild hybrids, SatFD207 and SatFH42 showed signals in two and 16 chromosomes, respectively (G–I). However, SatFH56 marked only two chromosomes in natural hybrids (I, J). In natural hybrids (G-J), signals from SatFD95 probes were observed in four chromosomes, among which in two of these signals colocalized with SatFH42 (indicated by arrows in H) and SatFH56 (indicated by arrows in J). Bars =10 µm.

**Figure 2.**
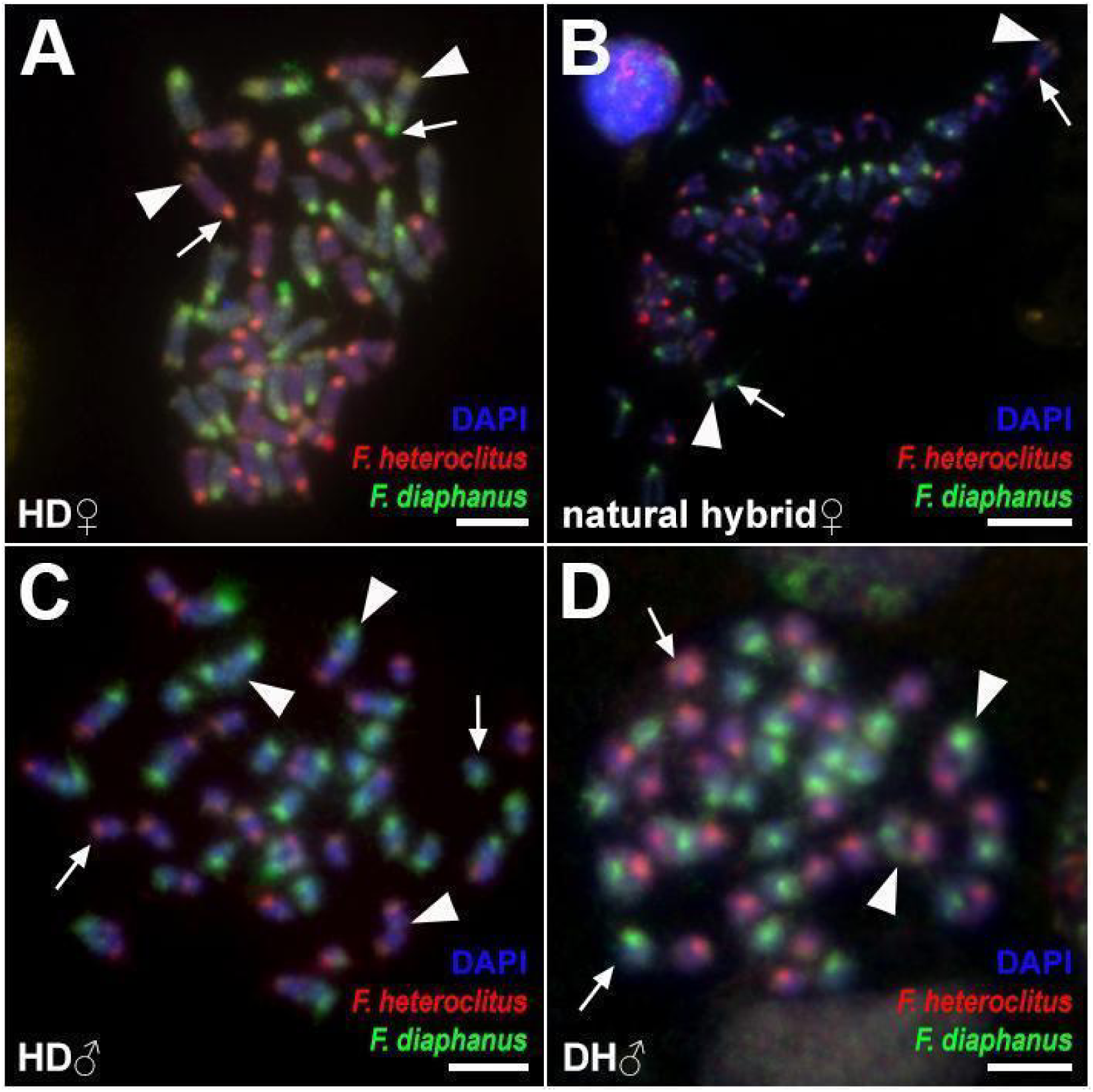
Comparative genomic hybridization reveals parental genome integrity in somatic metaphases (A, B) and meiotic pairing aberrations (C, D) in F1 lab-bred (A, C, D) and natural (B) *Fundulus* hybrids. Both parental genomes are clearly distinguishable in mitotic metaphases of F1 H×D hybrids (A) and natural (B) hybrid D×H females, showing the accumulation of species-specific repeats in pericentromeric regions (indicated by arrows) and subtelomeric regions of some chromosomes (indicated by arrowheads). Meiotic I metaphases from H×D (C) and D×H (D) hybrid males include univalents (pointed by arrows) and bivalents (pointed by arrowheads). Bivalents were observed between H and D, H and H, and D and D chromosomes. Red dye represents *F. heteroclitus* genomic DNA; green dye represents *F. diaphanus* genomic DNA. Bars = 10 µm.

We then identified eight species-specific satellite motifs: four were specific to *F. heteroclitus* (SatFH121, SatFH42, SatFH56 and SatFH213) and four others that were specific to *F. diaphanus* (SatFD150, SatFD233, SatFD95 and SatFD207) (Roussel et al. 2025). Fluorescent *In Situ* Hybridization (FISH) mapping of SatFH121, SatFH42, SatFH56, and SatFH213 on *F. heteroclitus* chromosomes revealed that only two produced strong signals (SatFH42 and SatFH56, Figure 1A). SatFH56 showed strong signals in the pericentromeric regions of two chromosome pairs and weaker signals in those of three additional pairs of chromosomes. SatFH42 exhibited bright signals in the centromeric regions of 12 chromosome pairs, weak signals in the pericentromeric regions of four pairs, and no detectable signal on the remaining eight pairs (Figure 1A). Dual-colour FISH with SatFH42 and SatFH56 revealed overlapping signals on one chromosome pair, while two additional pairs showed signals for SatFH56 only (Figure 1A). We did not observe sex specific differences in the localization of either satellite for *F. heteroclitus*. Mapping of SatFH42 and SatFH56 on *F. diaphanus* chromosomes showed no detectable signals (data not shown), confirming they were specific to *F. heteroclitus*.

FISH mapping of species-specific SatFD150, SatFD233, SatFD95, and SatFD207 on *F. diaphanus* chromosomes revealed clear signals only for SatFD95 and SatFD207 (Figure 1B). SatFD95 exhibited strong signals in the pericentromeric regions of two chromosome pairs and weak signals on three additional chromosome pairs. SatFD207 showed bright signals in the interstitial regions of two chromosome pairs, with an additional weak signal on the subtelomeric region of the p-arm on two pairs (Figure 1B). Dual-colour FISH revealed that SatFD95 and SatFD207 do not colocalize. We did not find sex specific differences in the localization of both satellites for *F. diaphanus*. We did not detect any signal for SatFD95 and SatFD207 repeats on *F. heteroclitus* chromosomes (data not shown), confirming these probes are specific to *F. diaphanus* (Figure 1B).

To assess parental genome integrity following hybridization, we mapped SatFH42, SatFH56, SatFD95, and SatFD207 onto the chromosomes of lab-bred F1 and wild asexual hybrids. The FISH mapping of these repeats revealed that the number of signals on chromosomes from lab-bred F1 fish (Figure 1C-F) was approximately half of that observed in the corresponding parental species, reflecting their hybrid chromosome composition. In asexually reproducing wild hybrids, SatFD233 and SatFH42 showed signal patterns similar to those observed in lab-bred F1 hybrids (Figure 1G–I). However, SatFH56 marked only two chromosomes in wild asexual hybrids (Figure 1I, J), as opposed to three in lab-bred F1 hybrids and six in *F. heteroclitu*s (Figure 1D, F). The SatFD95 probe marked four chromosomes in both F1 and natural hybrids. Surprisingly, two of these signals colocalized with SatFH42 (Figure 1H) and SatFH56 (Figure 1J) in asexually reproducing hybrids, a pattern we did not observe in lab-bred F1 hybrids. Thus, these data suggests a slightly different SatFD95 and SatFH56 distribution in natural hybrids compared to F1 hybrids and their sexual species.

### Hybrid males show dysfunctional gametogenesis while hybrid females have normal gonadal morphology

To investigate gametogenesis, we analyzed gonadal microanatomy using confocal microscopy. Germ cells were visualised by immunofluorescent staining of Vasa protein, which is highly expressed in germ cells and broadly used for their identification (Kobayashi et al. 2002; Begum et al. 2022). Germ cell types were distinguished based on nuclear morphology following DAPI staining, as previously reported in *Cobitis, Clarias, Hypseleotris,* and *Hexagrammos* (Dedukh et al. 2021; Majtánová et al. 2021; Dedukh et al. 2024; Dedukh et al. 2025).

In pure *F. diaphanus* (Figure 3A) and *F. heteroclitus* (data not shown) males, we detected distinct clusters of germ cells, including spermatocytes in pachytene and metaphase I, and large clusters of spermatids (haploid cells that will then undergo spermiogenesis to mature into sperm cells), as expected in fertile males. In addition, a few cells undergoing apoptosis were detected on gonadal tissue fragments (Supplementary Figure S1A). In natural (n=1) and lab-bred F1 D×H and H×D hybrid males (n=14) we detected gonocytes, pachytene cells, and large clusters of cells in metaphase I, but no spermatids (Figure 3B-D). These hybrid males also had clusters of cells with aberrant chromatin distribution, with many cells undergoing apoptosis (Supplementary Figure S1B-D). Tubulin staining, allowing for spindle visualization, revealed numerous spermatocytes in metaphase I with misaligned bivalents or univalents in hybrid males, suggesting arrest at this stage and subsequent accumulation (Figure 2C, D; 3D). Nevertheless, some metaphase I spermatocytes exhibited no misaligned univalents and appeared to segregate properly in hybrids from both cross types (Figure 3D). Altogether, these observations suggest that hybrid males from both cross types have decreased fertility and possibly complete infertility, consistent with gonad examination showing decreased testes size (data not shown, pers. obs., A.C.D.). Note that wild hybrid males are rarely found in Porters Lake (Fritz & Garside 1974; Dalziel et al. 2020; Hernandez Chavez & Turgeon 2007; Mérette et al. 2009); the single hybrid male we collected had gonads similar to lab-bred H×D and D×H hybrids, suggesting similarly reduced fertility in wild males (Figure 3E).

**Figure 3.**
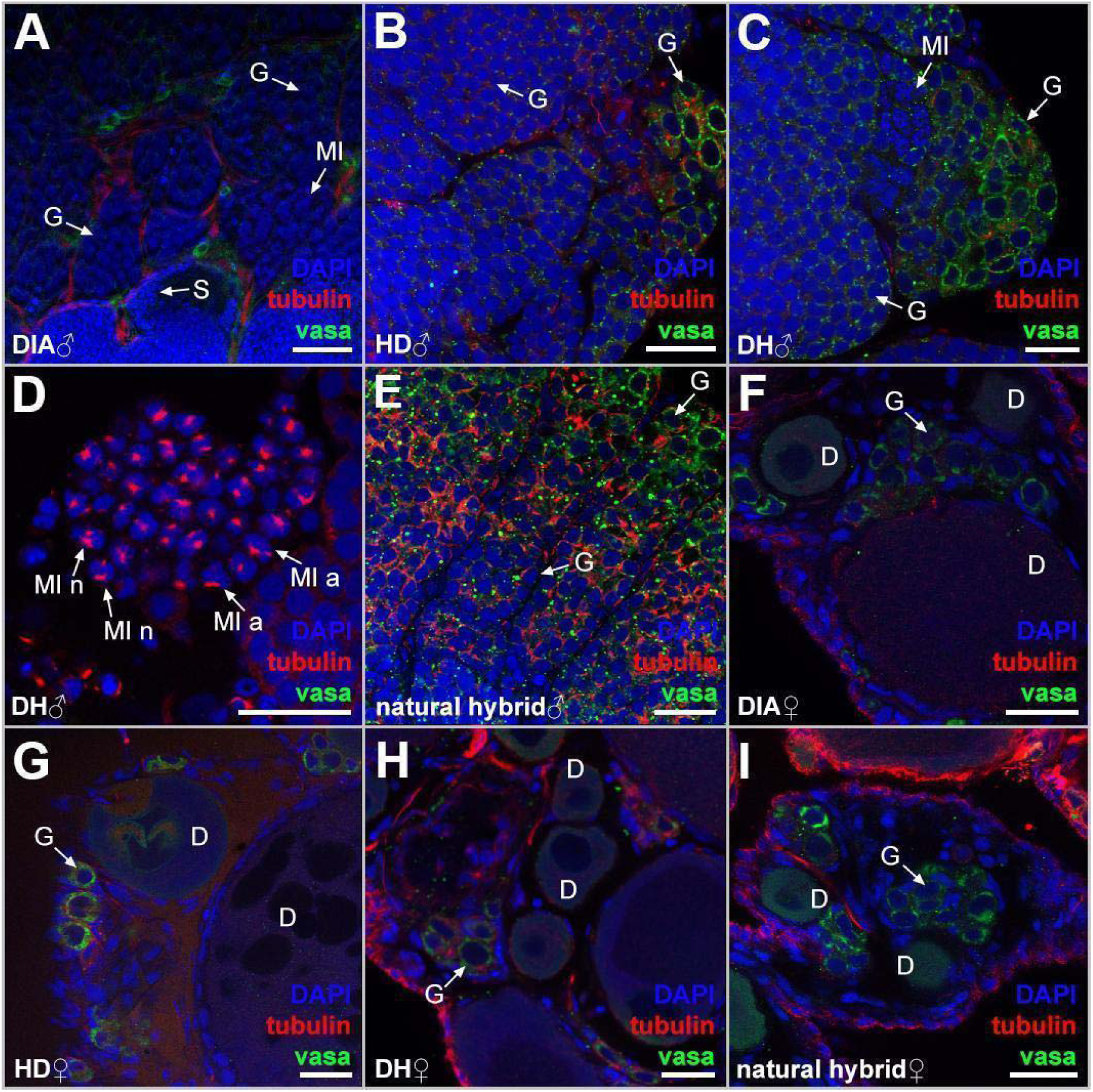
Gonadal microanatomy in sexual (A, E), F1 lab-bred hybrids (B-D, F-G) and natural hybrids (H). Whole-mount immunofluorescent staining with anti-Vasa antibodies (green) to visualise germ cells (G), tubulin (red) to visualise microtubules, and DAPI to stain chromatin (blue). (A) Sexual male *F. diaphanus*; (B) hybrid F1 lab-bred H×D male, (C, D) hybrid F1 lab-bred D×H male; (E) sexual female *F. diaphanus*; (F) hybrid F1 lab-bred H×D female, (G) hybrid lab-bred F1 D×H female ; (H) natural hybrid female (D×H). According to the morphology of nuclei, several germ cell line types can be determined: S, spermatids; G, germ cells; MI, metaphase plates during metaphase of first meiosis; MI n, metaphase plates with morphologically normal alignments of meiotic chromosomes in the equatorial pole and symmetrical spindle; MI a, metaphase plates with morphologically aberrant alignments of meiotic chromosomes in the equatorial pole and asymmetrical spindle; D, diplotene cells of meiotic division. Bars = 25 µm.

In *F. diaphanus* (Figure 3F) and *F. heteroclitus* (data not shown) females, we observed gonocytes and pachytene clusters located between pre-vitellogenic and vitellogenic oocytes, as expected for fertile females. In both H×D and D×H lab-bred F1 hybrid females, as well as in asexually reproducing wild D×H hybrid females, we observed all stages of germ cell development, including pre-vitellogenic and vitellogenic oocytes, suggesting hybrid females are fertile (Figure 3G-I). This is further supported by the production of healthy, live backcross progeny from H×D F1 females in laboratory conditions, although it remains unknown whether these progeny were actual backcrosses or clones (personal observations, A.C.D.). While a few apoptotic cells were detected in gonadal tissue fragments of *F. heteroclitus*, natural hybrids and H×D and D×H lab-bred F1 hybrid females, they are not predicted to substantially decrease fertility (Supplementary Figure S1E-H).

### Hybrid females from natural asexual lineages have premeiotic genome endoreplication

We investigated chromosomes in pachytene, a meiotic stage in which homologs should be completely synapsed, in wild fish, including pure *F. diaphanus* males (n=4) and females (n=5), pure *F. heteroclitus* males (n=4) and females (n=5), and D×H asexually reproducing females (n=5) (Supplementary Table 1). To confirm bivalent formation during the pachytene stage in parental species, we stained the axial (SYCP1) and lateral (SYCP3) elements of the synaptonemal complex. As expected from mitotic karyotypes, individuals from parental species of both sexes exhibited 24 bivalents (Figure 4A, Supplementary Figure S2A), each showing at least one crossover on pachytene spreads (Supplementary Figure S3A, B). Crossovers were visualised by α-MLH1 antibodies, which recognizes a subunit of the MLH1-MLH3 endonuclease responsible for Holliday junction resolution and crossover formation, on pachytene chromosomes (Moens et al., 2002). We also visualized DNA double-strand breaks (DSB) using immunodetection of RAD51, a key protein involved in homology search and DNA double-strand break (DSB) repair during meiosis (reviewed in Ito et al. (2024)). During early pachytene, we observed intense RAD51 staining along the synaptonemal complexes, consistent with numerous DSB being induced along each bivalent in metazoans, including fish (reviewed in de Massy (2013) and Nath et al. (2022)). This intense staining was followed by a progressive decrease in the number of RAD51 foci through mid- and late pachytene, resulting from DSB repair, as expected for normal meiotic progression in fertile animals (Supplementary Figure S2C, D).

**Figure 4.**
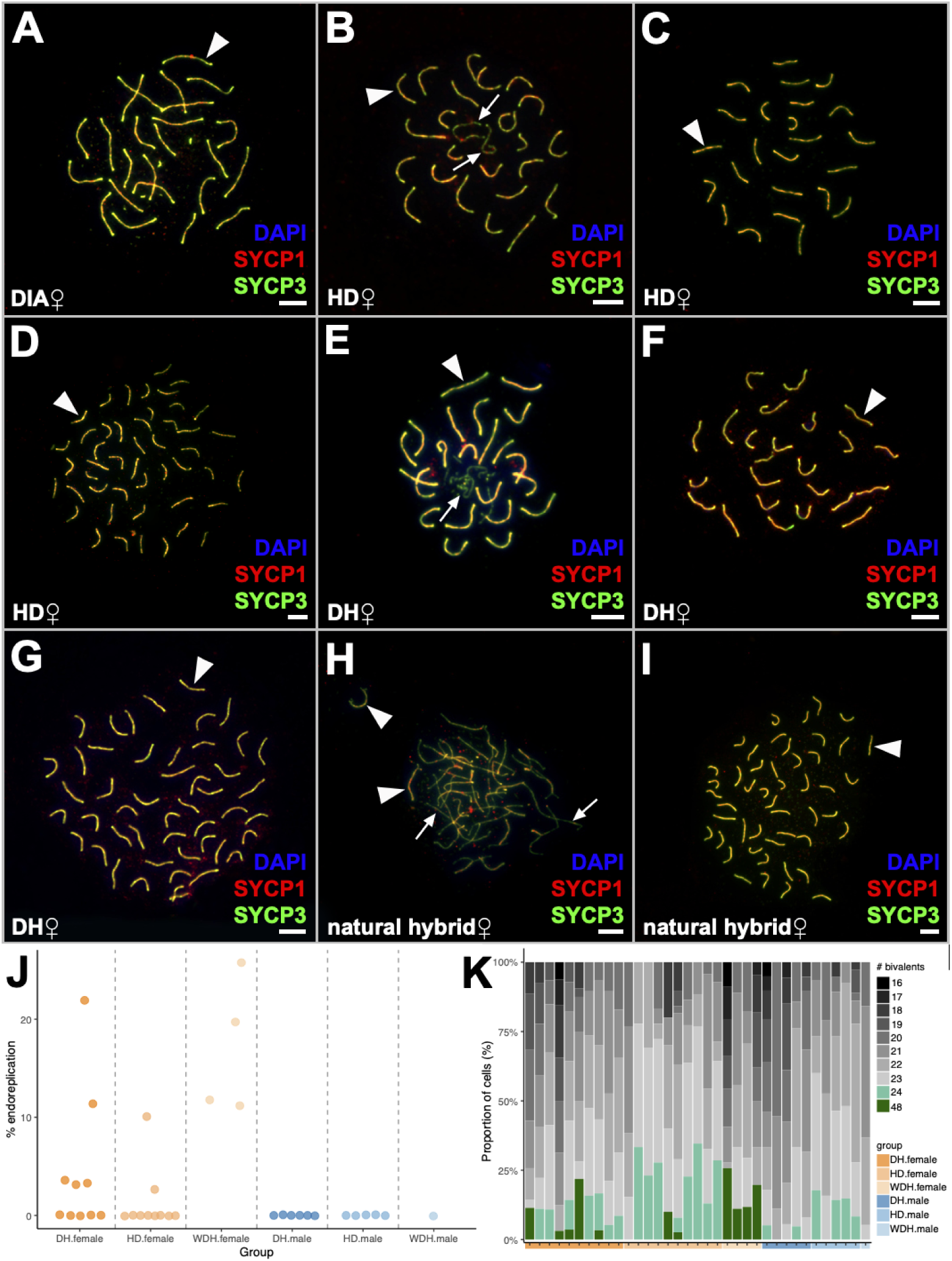
The analysis of pairing in pachytene oocytes of sexual individuals (A), F1 lab-bred hybrids (B-G), and natural hybrids (H-I). Synaptonemal complexes were visualized using immunostaining of lateral (SYCP3 protein, green) and central (SYCP1 protein, red) components; chromatin is visualized by DAPI staining (blue). SYCP3 and SYCP1 proteins (indicated by arrowheads) accumulate on bivalents, while SYCP3 protein (indicated by arrows) accumulates only on univalents. Scale bar = 10lµm. J. Proportion of cells showing endoreplication for each group of fish. K. Proportion of cells showing a given number of properly aligned bivalents for each group.

We further examined pachytene pairing in four natural D×H hybrid females and one rare D×H hybrid male. In females, among 193 pachytene oocytes, 78% maintained diploidy, while 22% were tetraploid (11.1-25.7% of the cells across 4 individuals). Among “normal” diploid pachytene oocytes from these hybrid females, we observed normal pairing for an average of 21 ± 2 bivalents, but an average of 6 ± 4 chromosomes remained as univalents (Figure 4H). Crossovers were observed on bivalents, but were absent on univalents or unpaired chromosomal regions (Supplementary Figure S3). DSBs were predominantly found on univalents or unpaired chromosomal regions, whereas bivalents exhibited only a few signals (Supplementary Figure S3). Unlike results from hybrid diploid oocytes, we found no aberrant pairing or univalents in tetraploid oocytes with 48 bivalents (Figure 4I). Given that these oocytes had a doubled chromosome number (4n = 96), they likely emerged through premeiotic genome endoreplication. At least one crossover signal was detected per bivalent, confirming successful pairing and suggesting normal DNA repair and crossover (Supplementary Figure S3M, N). We were unfortunately unable to detect DSBs in tetraploid pachytene cells for technical reasons.

### Cross direction affects endoreplication and chromosome synapsis in lab-bred F1 hybrid females

We further analyzed gametogenesis in reciprocal lab-bred F1 hybrid females (D×H: n = 10; H×D: n = 10). Among the ten D×H hybrid females, five had two distinct populations of pachytene oocytes: one population with 2n = 48 chromosomes (i.e., diploid) and one with 4n = 96 chromosomes (i.e., tetraploid) (Figure 4E-G). In the remaining five D×H females, only diploid oocytes were observed (Figure 4E, F, Supplementary Figure S3I-L). Among the ten H×D hybrid females, only two individuals exhibited both diploid and tetraploid oocytes (Figure 4B-D). Similarly to natural D×H asexual hybrids, tetraploid oocytes in both H×D and D×H hybrid females contained 48 fully paired bivalents (Figure 4D, G). Our statistical analysis of endoreplication frequency showed that genotype had a significant effect on frequency of endoreplication in females (Figure 4J; H_(2)_ = 10.954, p = 0.004). In particular, there was a significantly higher frequency of endoreplication in wild D×H hybrid females thanlab-bred F1 D×H (z = −2.314, p = 0.021) and F1 H×D hybrid females (z = −3.132, p = 0.002), but no significant difference between lab-bred F1 D×H hybrid females and lab-bred F1 H×D hybrid females (z = −1.508, p = 0.132).

Among diploid oocytes, the average number of correctly paired chromosomes per cell differed by genotype among female hybrids (Figure 4K; F_(2,20)_ = 7.125, p = 0.005). The frequency of proper pairing was significantly lower in wild D×H hybrid females than lab-bred F1 H×D hybrid females (p = 0.006), but not lab-bred F1 D×H hybrid females (p = 0.175). No differences in correctly paired chromosomes per cell were detected between reciprocal lab-bred F1 D×H hybrid females and lab-bred F1 H×D hybrid females (p = 0.092). In lab-bred F1 H×D hybrids, the number of fully paired bivalents ranged from 16 to 24 (average 21.3 ±0.3), while in D×H hybrids, the range was 18 to 24 (average 22.2 ±0.2).

The proportion of diploid cells in which all pairs were properly aligned also varied by genotype among female hybrids (Figure 4K; F_(2,20)_ = 6.666, p = 0.006). Lab-bred F1 H×D hybrid females exhibited a significantly higher proportion of properly aligned diploid cells than wild or lab-bred D×H hybrid females (p = 0.009 and p = 0.047, respectively). There was no significant difference in the proportion of properly aligned pairs between the wild D×H hybrids and the lab-bred F1 D×H hybrids (p = 0.569). Similar to wild asexual D×H hybrids, crossovers were detected on bivalents, but not on univalents (Supplementary Figure S2E, I). RAD51 loci intensely decorated univalents and unpaired chromosomal regions, while only a few signals were detected on bivalents (Supplementary Figure S3F, J). These results suggest that oocytes with unsynapsed chromosomes and unrepaired DSBs cannot progress beyond meiosis and form gametes, likely resulting in decreased fertility in lab-bred F1 hybrid females from both cross-directions.

### Lab-bred F1 and natural hybrid males do not exhibit premeiotic genome endoreplication

We analyzed gametogenesis in a single wild D×H hybrid male and in lab-bred F1 hybrid males from H×D (n = 5) or D×H (n = 6) crosses (Supplementary Figure S2). In contrast to hybrid females, we did not observe any pachytene cells with a duplicated genome, and all spermatocytes retained their initial ploidy (Supplementary Figure S2F, G). In both wild and lab-bred F1 hybrid males, spermatocytes exhibited aberrant pairing, forming an average of 20 ± 2 bivalents with 6 ± 4 chromosomes remaining unpaired (Supplementary Figure S2F, G). However, in some lab-bred hybrids, we additionally observed rare pachytene spermatocytes with complete synapsis of all 48 chromosomes into 24 bivalents in both cross types. Crossovers were detected on bivalents, but not on univalents (MLH1 staining, Supplementary Figure S3G, K), while RAD51 loci were observed on univalents mostly, suggesting delayed DNA DSB reparation (Supplementary Figure S3H, L). We conclude that lab-bred and natural hybrid males likely have very low fertility due to aberrant chromosomal pairing in the majority of spermatocytes, and predict that some individuals are completely sterile.

## Discussion

Natural, all-female, clonal *F. diaphanus* × *F. heteroclitus* lineages have repeatedly arisen in some regions where these species overlap, but the mechanisms underlying asexual reproduction remained unknown. We found that: 1) all wild D×H asexual females can produce diploid eggs via premeiotic endoduplication, 2) the gametogenic alterations found in these wild clonal lineages are found in some lab-bred reciprocal F1 females, but at a reduced rate, and 3) wild and lab-bred reciprocal F1 male hybrids show no evidence for endoduplication and while rare spermatocytes do show proper pairing and synapsis, hybrid males are likely infertile.

### Wild asexual and lab-bred hybrid females produce two types of oocytes

To determine the mechanisms leading to asexual reproduction in female *Fundulus* spp. hybrids, we analyzed gametogenesis in natural D×H asexual hybrid females, lab-bred reciprocal F1 hybrids and wild and lab-bred pure parental species*, F. diaphanus* and *F. heteroclitus*. By observing the synaptonemal complex (SC) forming between homologs in prophase I, we observed two types of oocytes in natural (D×H) and both types of lab-bred F1 hybrid females: (1) diploid oocytes (2n = 48), and (2) tetraploid oocytes (4n = 96) with a doubled genome resulting from premeiotic endoreplication. Premeiotic endoreplication is a widespread gametogenic alteration, allowing identical chromosome copies to pair during meiosis instead of orthologs (e.g., Lutes et al. 2010; Kuroda et al. 2018; Dedukh et al. 2020). Crossovers were detected on all bivalents, but, in contrast to sexual species, they occur between identical chromosomal copies, resulting in the production of clonal gametes. These tetraploid oocytes can further progress through meiosis and form diploid gametes, which are genetically identical to the mother. This mechanism efficiently resolves pairing challenges of divergent orthologous chromosomes during prophase I in hybrids, and has been reported in other clonal fishes, amphibians, and reptiles (Macgregor & Uzzell 1964; Cimino 1972; Stöck et al. 2002; Lutes et al. 2010; Kuroda et al. 2018; Dedukh et al. 2020; Dedukh, Altmanová, et al. 2022; Dedukh et al. 2024).

In contrast to wild hybrid females, not all lab-bred F1 hybrid females produced tetraploid oocytes and those that did produced a lower proportion than natural asexual females. Of the two reciprocal crosses, a higher percentage of lab-bred D×H hybrids (the cross direction that predominates in the wild) produced tetraploid oocytes, at a higher proportion, than H×D hybrids, although the difference was not statistically significant. Thus, premeiotic endoreplication leading to diploid clonal gamete formation occurs in lab-bred *Fundulus* F1 hybrids, as reported in *Cobitis*, *Oryzias*, and clariid fishes (Shimizu et al. 2000; Choleva et al. 2012; Dedukh et al. 2021; Marta et al. 2023; Dedukh et al. 2024). Together, these data suggest that new clonal lineages may still arise in nature when *F. diaphanus* and *F. heteroclitus* hybridize (Dalziel et al. 2020).

Endoreplication can occur through three main pathways: 1) cell fusion, where two gonial cells merge; 2) endomitosis, where DNA is replicated followed by chromosome condensation, but mitosis or chromosome segregation fails; and 3) endoreduplication (or endocycling), which involves repeated rounds of DNA synthesis and mitosis bypass. While the exact mechanism underlying endoreplication in asexual *Fundulus* spp. hybrids remain unclear, it is unlikely that cell fusion occurs, as we did not detect binucleated oogonia by confocal microscopy, unlike in laboratory strains of asexual *Carassius* hybrids, which exhibit cell fusion events (Wang et al. 2016). In *Cobitis* hybrids, endoreplication likely occurs shortly before meiosis, as no clusters of oogonia or pachytene oocytes with duplicated genomes have been detected (Dedukh et al. 2021). Supporting a mechanistic link between endomitosis and endoreplication, Rotelli et al. (2019) showed that stress-induced endoreplication in *Drosophila* somatic cells involves the same cell cycle regulators for both endomitosis and endoreduplication—Cyclin A/CDK1 and Aurora B kinase. Specifically, complete inhibition of these regulators triggers endoreduplication, whereas partial knockdown results in endomitosis. Altogether, these findings suggest that the observed tetraploidy in *Fundulus* female hybrid oocytes is unlikely to result from cell fusion, and may instead reflect a regulatory shift toward endoreduplication or endomitosis.

In most known asexual hybrid vertebrates, premeiotic genome endoreplication is a rare event in germ cells (Shimizu et al. 2000; Newton et al. 2016; Dedukh et al. 2021; Marta et al. 2023; Dedukh et al. 2024; Dedukh, Altmanová, et al. 2022; Spangenberg et al. 2025). Consistently, the majority of pachytene oocytes (78% in 4 wild females and 97% in lab-bred F1 hybrid females) maintained their normal ploidy. In diploid oocytes with unduplicated genomes, the average number of properly paired chromosomes was higher in lab-bred H×D than natural D×H hybrids, but did not significantly differ between lab-bred D×H and wild D×H hybrids. There was a trend towards a difference between lab-bred H×D and lab-bred D×H hybrids, but it was not significant. It is worth noting that there was a significant difference in the proportion of diploid cells with properly aligned chromosomes with lab-bred H×D hybrids having significantly more cells with full pairing than wild and lab-bred D×H hybrids. All together, these results suggest an effect of the cross type on the probability of successfully completing meiosis I for diploid oocytes with an unduplicated genome.

Previously, no significant differences in chromosomal pairing were observed between pure *Cobitis* species and reciprocal F1 crosses (Marta et al. 2023). Nevertheless, some crosses exhibit a drastic bias in fertility, showing clonal gametogenesis in one direction and sterility in the other. Although direction-specific biases have been detected in other organisms, and these are often explained by the influence of sex chromosomes, this phenomenon still requires further investigation in species with homomorphic sex chromosomes (Stöck et al. 2021). The analysis of gonadal morphology revealed normally looking ovaries with various germ cell lines, indicating their potential fertility. The presence of oocytes with 24 bivalents suggests that some lab-bred F1 females can also form recombined gametes in addition to diploid clonal ones, matching findings of rare backcross hybrids in Porters Lake (Hernandez-Chavez and Turgeon, 2007). Simultaneous formation of meiotic and clonal gametes has also been detected in lab-bred F1 *Cobitis* hybrids (Janko et al., 2018; Marta et al. 2023).

The high number of diploid oocytes with univalents and unrepaired DNA DSB likely implies a substantial fertility reduction. However, such a decrease in fertility may facilitate the long-term stabilization of sexual–asexual coexistence, because gynogenetic lineages depend on the sperm of sexual species to activate development (Leung & Angers 2018). Generally, the reduction in fecundity of asexuals is less pronounced than in parental species, especially given the high egg production typical for many teleosts. For example, triploid *C. elongatoides* × *C. taenia* females exhibit a ∼50% reduction in fecundity compared to *C. taenia* (Juchno & Boroń 2006, 2010), which is far less than expected from the extensive loss of pachytene oocytes (Dedukh et al. 2021). Moreover, the parthenogenetic lizard *Aspidoscelis neomexicana* also has a similar fecundity to that of its sexual ancestors, despite most pachytene oocytes not progressing beyond pachytene (Newton et al. 2016). These data suggest that fecundity may not be dramatically reduced in hybrid *Fundulus* females. Instead, fertility appears to be maintained through oocytes with endoreplicated genomes, which continue to accumulate nutrients during the diplotene stage and ultimately give rise to progeny. Although the loss of non-duplicated pachytene cells may be problematic, it is not critical for fertility, as resource investment is concentrated in later stages of oocyte development.

Taken together, lab-bred hybrids exhibit significantly lower rates of endoreplication than natural clones, suggesting interclonal selection might favor hybrids with higher rates of endoreplication. Following their establishment, it is possible some clones may evolve to further increase endoreplication rates. Supporting the persistence of asexual *Fundulus* clones for at least ∼30 generations, we observed slightly rearranged satellite repeats compared to parental species and lab-bred reciprocal F1 hybrids. A similar pattern has been reported in *Cobitis* hybrids, where hybridization between sexual species produces clonal gametes via endoreplication (Dedukh et al. 2021; Marta et al. 2023). Nevertheless, only a limited number of clonal lineages persist in natural populations, and these are characterized by elevated rates of premeiotic genome duplication and higher fecundity (Janko et al. 2005).

### Male hybrids do not appear to be fertile

In contrast to hybrid females, no males displayed genome endoreplication (i.e., X males from reciprocal crosses and a single wild hybrid D×H male). Among pachytene spermatocytes with abnormal pairing, we found an average of 16 to 22 of 24 bivalents, and 4 to 16 corresponding univalents. Interestingly, some spermatocytes with aberrant pairing progressed beyond pachytene checkpoints and entered meiosis. Similarly to pachytene spermatocytes, we detected cells with both H and D univalents but also HH, DD, and HD bivalents. On gonadal tissue fragments, we rarely observed spermatids and sperm cells. In metaphase I, we observed aberrant chromosome attachments to the spindle, possibly due to univalent formation. Hence, we suggest that a large proportion of spermatocytes are unable to progress beyond metaphase I, likely decreasing hybrid male fertility. Decreased fertility or complete sterility in hybrids is consistent with the reduced size of their gonads compared to parental species.

The mechanisms behind this sex-specific susceptibility to genome endoreplication remain unclear, and could be triggered by: 1) sex determination, i.e., the genetic or environmental cue establishing an individual as a male or female, or 2) sexual differentiation, i.e., the process by which sexual traits develop. For example, work on *Cobitis* hybrids showed that spermatogonia from male hybrids transplanted into the ovaries of sexual females gained the ability to endoreplicate their genomes (Tichopád et al. 2022). This result suggests that sexual differentiation, through the gonadal environment, rather than sex determining cues (genetic or environmental), controls genome endoreplication. On the contrary, in *Misgurnus* loaches, endoreplication occurs in genetic female hybrids that are sex-reversed into males by hormonal treatment (Yoshikawa et al. 2007), indicating that sex determination is a key factor influencing genome endoreplication. A better understanding of the genetic, molecular and cellular mechanisms underlying sex-biased genome endoreplication in various systems is required to understand this natural phenomenon.

Several studies have shown that asexual vertebrate hybrids tend to arise from genetically divergent species, often with homomorphic sex chromosomes (Janko et al. 2018; Stöck et al. 2021). In *Cobitis* hybrids, karyotype similarity appears to have contributed to the emergence of asexual gametogenesis (Marta et al. 2023). *F. diaphanus* and *F. heteroclitus diverged* 15-25 million years ago, have similar karyotypes (2n = 48) (Ghedotti & Davis 2017) with homomorphic sex chromosomes. This suggests that studied hybrids inherit two key traits promoting asexuality: high genetic divergence and homomorphic sex chromosomes, matching data from known systems (Janko et al. 2018; Stöck et al. 2021; Marta et al. 2023).

### Wild asexually reproducing D×H females as quasi “frozen F1” hybrids

We found no rearrangements or genomic introgression between *F. diaphanus* and *F. heteroclitus* chromosome sets in clonally reproducing hybrids by CGH, suggesting that parental genomes remain mostly intact. These data match observations in other asexually reproducing vertebrate hybrid lineages displaying predominantly “frozen” parental chromosome sets over many generations (Majtánová et al. 2016, 2021; Zaleśna et al. 2011; Dudzik et al. 2023).

However, loss-of-heterozygosity events and differences in repetitive DNA distribution can occur over several generations of clonal genome transmission in asexual hybrids (Warren et al. 2018; Jaron et al. 2021; Janko et al. 2021; Vukić et al. 2025). Indeed, we detected subtle differences in repetitive DNA content between parental species, lab-bred F1 hybrids, and wild asexually reproducing hybrid females. Satellite FISH probes specific to *F. diaphanus* (SatFD233) and *F. heteroclitus* (SatFH42) showed similar patterns in the pure individuals and wild and F1 lab-bred hybrids. However, the *F. heteroclitus* specific (SatFH56) probe produced fewer signals in wild asexually reproducing hybrids than expected, suggesting a possible loss-of-heterozygosity event. In addition, the *F. diaphanus* specific (SatFD95) probe colocalized with the *F. heteroclitus* specific SatFH56 and SatFH42 probes in natural asexually reproducing hybrids – a pattern absent in reciprocal lab-bred F1 hybrids and parental species. This finding points to two possibilities: satellite repeats evolved independently during clonal propagation, or the ancestral genomic state of the parental species from the original hybridization event was not captured. Indeed, clonal and hemiclonal reproduction can preserve genomes of extinct species, enabling their “ghost” or “zombie” persistence in hybrids (Alves et al. 2001; Dubey & Dufresnes 2017; Unmack et al. 2019; Vukić et al. 2025). Moreover, without recombination, clonally transmitted genomes can undergo mutation accumulation and structural rearrangements, evolving independently in contrast to sexually transmitted genomes (i.e., the Meselson effect) (Vorburger 2001; Gromicho et al. 2006; Manaresi et al. 1992; Tinti & Scali 1992; Scali et al. 2020; Öztoprak et al. 2025). At present, we cannot rule out any of these possibilities.

## Acknowledgements

We would like to thank Abby Brouwer, Isadora Schumann Munhoz, Sarah Young Veenstra, and Julia McIsaac for their assistance with fish crosses and husbandry at Saint Mary’s University. We thank Claude Power for his help with field sampling, and Annabelle Fournier for her assistance with sample processing. We also thank Caila Henderson Lebans for her assistance with aquatic facilities management. We thank members of the Saint Mary’s University and Universite de Moncton Animal Care Committees for their work to ensure animal care met ethical standards.

## Author contribution

DD: Conceptualization; Funding acquisition; Investigation; Supervision; Writing—original draft; Writing—review & editing.

KO: Investigation; ZM: Investigation; PO: Investigation; AJR: Investigation;

MEH: Formal analysis; Writing—review & editing. KJ: Funding acquisition;

ACD: Conceptualization; Funding acquisition; Investigation; Supervision; Writing—original draft; Writing—review & editing.

AMDC: Conceptualization; Funding acquisition; Investigation; Supervision; Writing—original draft; Writing—review & editing.

