## Supplementary material for "Sex-specific modifications of gametogenesis in natural and lab-bred *Fundulus* spp. hybrids"

**Supplementary Figures**

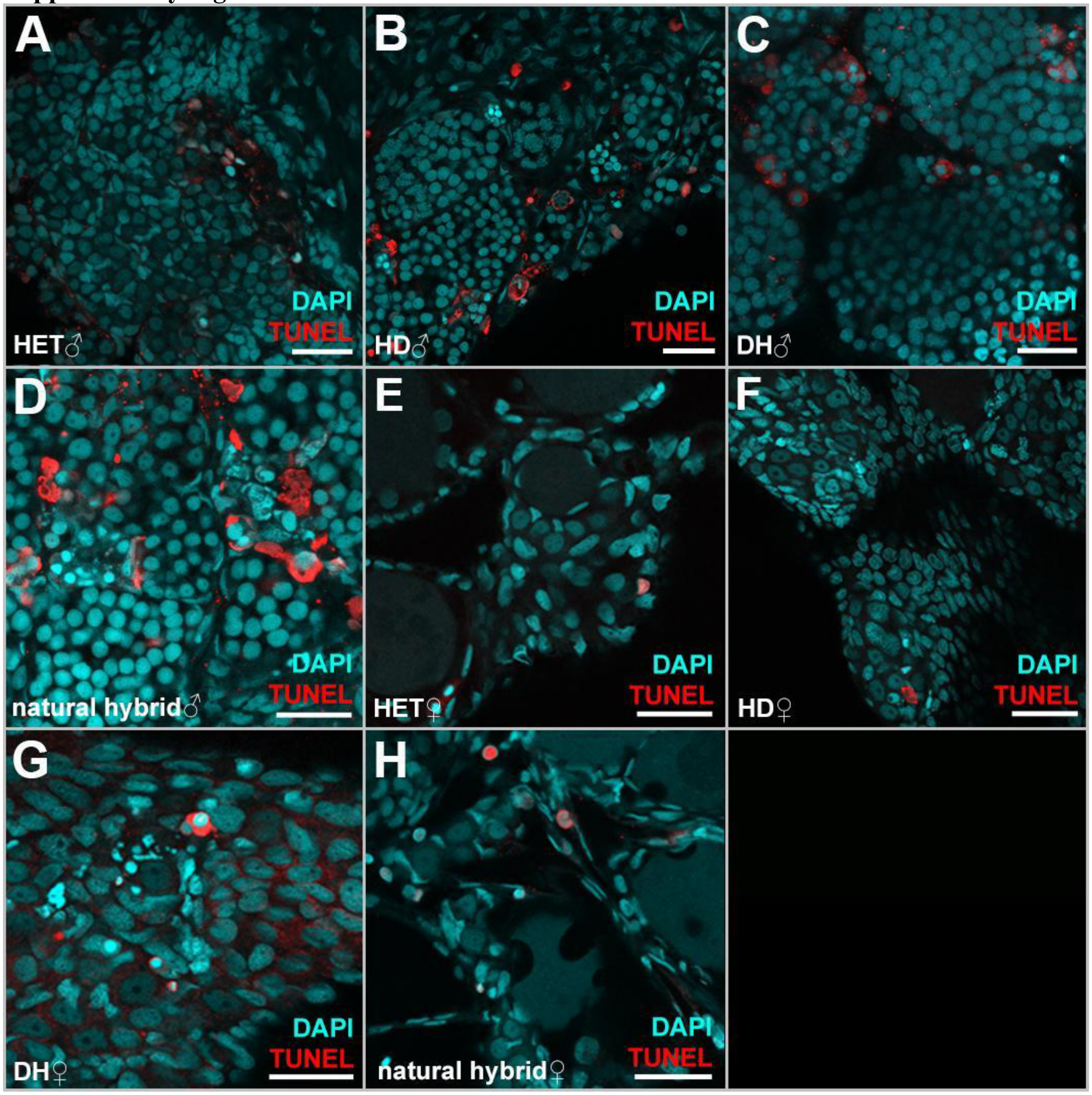


**Supplementary Figure S1. Detection of apoptosis in gonadal tissue fragments in sexual (A, E), F1 hybrids (B, C, F-G) and natural hybrids (D, H).** TUNEL assay (red) visualised DNA degradation and DAPI stained chromatin (cyan). (A) Sexual male *F. heteroclitus*; (B) hybrid F1 lab-bredH×D male , (C) hybrid F1 lab-bred D×H male; (E) sexual female *F. heteroclitus*; (F) hybrid F1 lab-bred H×D female, (G) hybrid  F1 lab-bred D×H female ; (H) natural hybrid female. Bars = 25 μm.

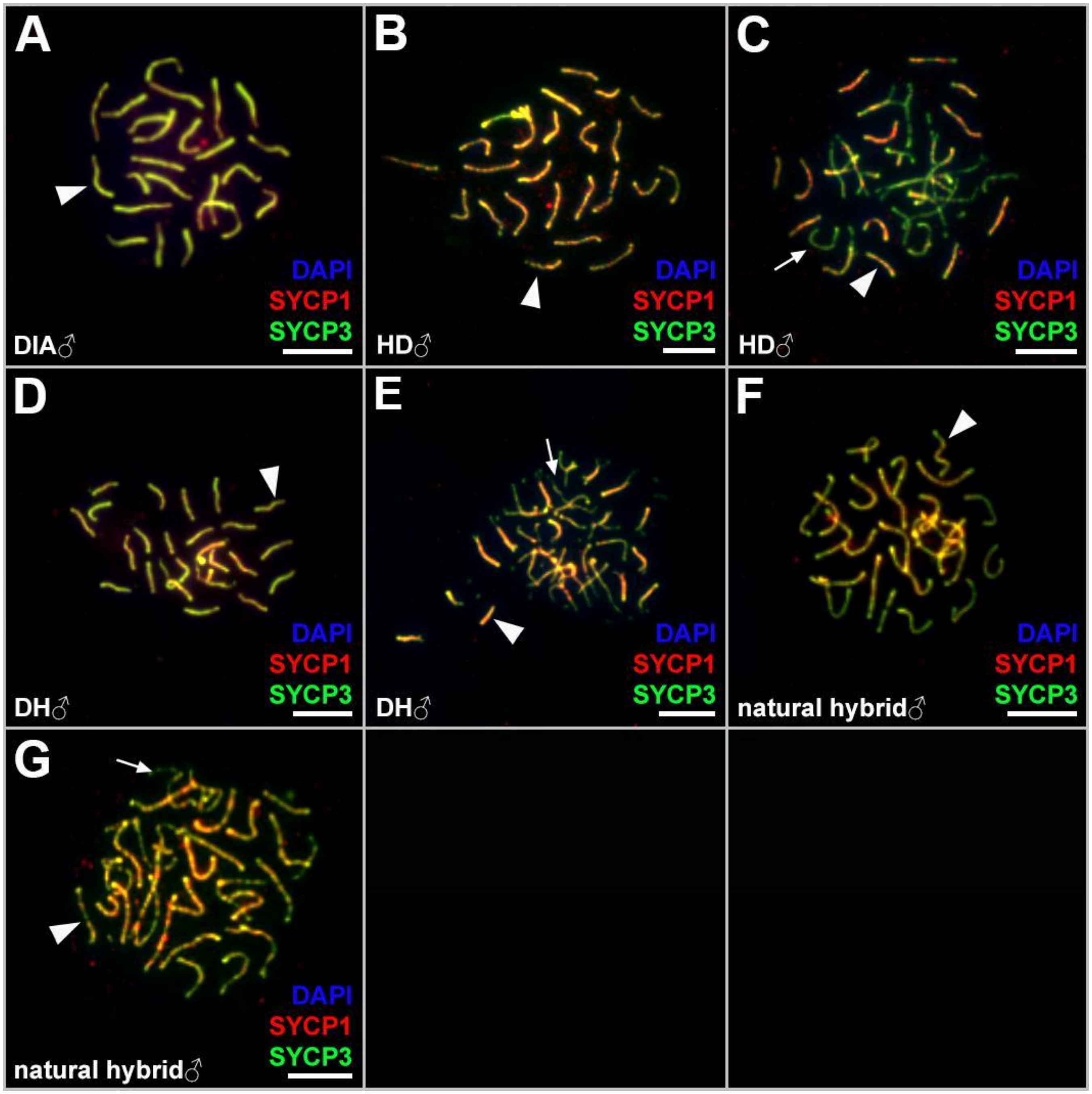


**Supplementary Figure S2. The analysis of pairing in pachytene spermatocytes of sexual individuals (A), F1 lab-bred hybrids (C-E) and natural hybrids (F-G).** Synaptonemal complexes were visualized by SYCP3 (green, lateral element) and SYCP1 (red, central element) immunostaining; chromatin was stained with DAPI (blue). Bivalents (arrowheads) show SYCP3 and SYCP1 signals, while univalents (arrows) show SYCP3 only. Scale bars = 10 µm.

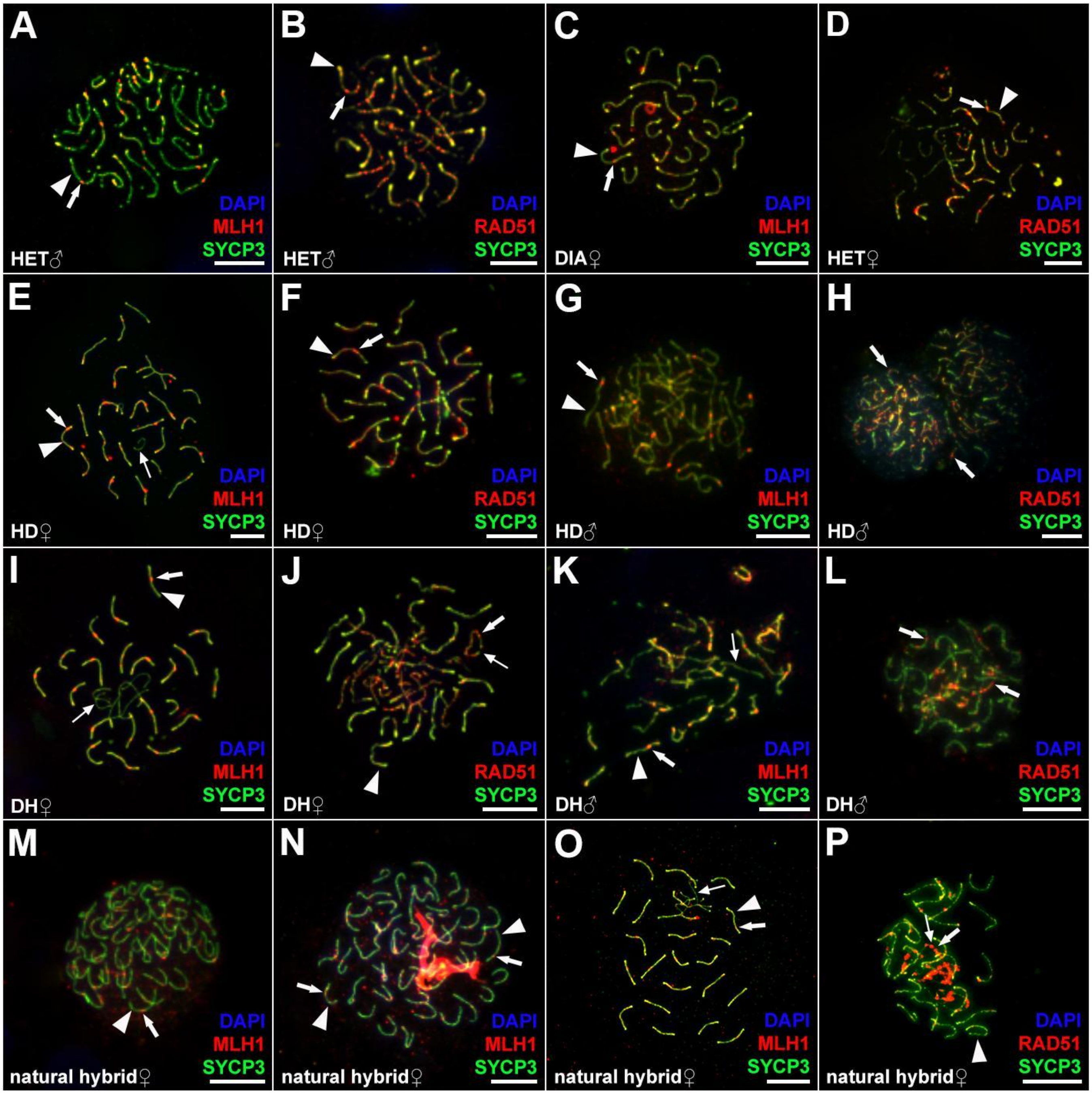


**Supplementary Figure S3. The analysis of recombination and double-strand brake formation in pachytene meiocytes of sexual individuals (A-D), lab-bred F1 hybrids (E-L) and natural hybrids (M-P).** Synaptonemal complexes were visualized by SYCP3 (green, lateral element); recombination loci were identified using anti-MLH1 antibodies (thick arrows in A, C, E, G, I, K, M-O); double-strand breaks were visualized by anti-Rad51 antibodies (thick arrows in B, D, F, H, J, L, P); chromatin was stained with DAPI (blue). Bivalents (arrowheads) were indicated by arrowheads, univalents by thin arrows. Scale bars = 10 μm.

**Supplementary Materials 1. Monomer sequence of each satellite repeat with the indication of primers and sequences used as oligoprobes.**

**SatFH121**
TTTTAGTTTGGGTTCAGGGTTAGGTTTAGGCATGGGGTGAGGGTGGGGTTAGGGTAAGGGTC
AGATACAGGGGTGGGGTTAGGAAAGGGAGACCCAGTGACCGCCGCTTGACCAGAGGCACG
GGCAGGTTAGGGAACCAAAGATATGACAGTTCCCCCCGAAGGGGGGAACTGGAATGCACTG
GCCATCATGGTGTCGGTCAGCCACCAAGTCTCCCATCCCCAGTCGCCTTGGCTATCTAATAAT
AATAAAGTTTTATCATGTCAAGTCCGGTGTTGTGTCGTCTCTTGTCTGGAGTTCTTCTGTTTTT
TGTAGTGTGCTTATAAGCTTTTCTTTGTCTGTCTATTTCCGTTCTTTTCTGTTTCTTTTATATGT
GCCTTTGTTGGCTTTTCTGTTTCTTGTTGTCTGTTTTTTTCTGTTTCTCTAATTTCTGCCGTTCTT
GGCTTTTCTGTTCCTCGTTTTCAATTACGGACGGCCTTGTCGACGACGTTTCGACGTAATTGT
GTCTTCTTCAGGTCTTGTAGCGTCCGTGTCTTCTGTGTAAATGTGTATTATTTCTCCTTTTATA
GGTTTTACAGGTAAGTATTTTCTTTTCCAGGTAAGTATTTCCGATTTCTTTTCAGGTTTGTGTT
TCTTCCAGGTAAGTATTTTTGCTTTTCAGGTTTTGGGTTTTTTTCGCGGTTCTAGGTAGGTATA
TTTATGTATTTTTTTGGTATTTTTTCCTTTTTCTGTTTTTTTCTGTTTTTTTCTATATGCTTTCCT
TAGCTTTTCTGTTTCTTTACCCCTCCAATAAAATGAGTTTAATTTGTTTCTTAATTCAACGCTT
TTCTGAGCTTTTCTGTTTTTTGATCCTTTCCCCTCTGTGTCCAATTTGAATTTGAAAAATCCCT
CGTCCTCCTGTTCCTTATATATCTCTTAATTTTAGATTTAATTTATCCCAAAATTTTCCATTTA
GTAATTCCTTATTTTAGGTTTGTATTTAGGTTTGTAAGTGGGATTGGGGTTAGGAAATAGGGA
ATTAGGGTTAGCTAGGGGTCAATAGGGGTCAGGATATGGAATGGGGTAAGGGTCGGATTAC
TTGCAGCCGCCAGAGAGACAGACCCAACCGTCCGTTTGCTGACGGGAATCCCAGCCATTGG
GCTTACTCTGCGGAAAACGTACCTCGCTCTGTGCGGAATTCTGTGGGGTCTTAGTCCTCTCAT
TGCCTCTCCCTGGGTCGCGGGATGCCCAATATCTCTGTCGCTACTAGGTCCCCTGGGCTGGG
ATCCCGTAGCTCCTGGAGGTGGGCCTGCCTCTCTCGCTGTCTTTCGAATCCGGGGTGGGGTA
GGGTTCTCTAATTTCTGTCTCTGTTTCTTATATATGGCACCCCCTGTTCTGTTCTTCTCTCTGT
CTTTCTTAATTTAGATAATTTTCTCTTCTTTGTTTCTCTAATTAGTTTAGGCACCCCCTATTTCC
TTTCTCTCTTTCTTTCTTTCCCTATTTAAACATCTTTTTTTTGA

**Primers:**
Forward: 5` TGTAGCGTCCGTGTCTTCTG 3`
Reverse: 5` TCCGCAGAGTAAGCCCAATG 3`
**Oligoprobe:**
5` GCAGCCGCCAGAGAGACAGACCCAACCGTCCGTTTGC 3` biotin or digoxigenin

**SatFH42**
GCGGTTTGATATCTCCAAGTTAAGTAGAATCTTTACTGGTGTCTCTTTGGGCCTGGTGGGGCT
CCATGAAGCCTAAATCCTTGAGCTAGGGTTGGGATCTTTGGTCAGGCTCATCCCAGAGGGCT
GATTTTCACAAATGTCATTCAGGGGGTGACAATGAGTGGATTCTGTGAGTTTGGTGTCGCTG
GAGCGTTTCGTCTTGGCATGACAGGGGCAGGAAGGGGTGGAAAAATGACCAAAAATGAAAC
TTTGACACCTCATAACTCTGAGACGTTTGGTTTAATCTGGCCCAAATTTGCTATGCAGCTTCC
TGTGGTAAGGTGCTACAAACCTGGAAGAGCCTATCTCTCAAAGACCTTCCATCTGTTAGTTG
TTTTTGGACCGGTTATTAGGTTTGACGGCACAGCACGGCTTTCTCTCGGCCCCTGGGGCAGAT
AGGGACCCCAAATTACCAGGGCCTGCTCTACTCACACCCTAGAAACCGT
**Primers**:
Forward: 5' TCTGTGAGTTTGGTGTCGCT 3`
Reverse: 5` GTGCTGTGCCGTCAAACCTA 3`
**Oligoprobe**:
5` GAGACGTTTGGTTTAATCTGGCCCAAATTTGCTATGC 3` biotin or digoxigenin

**SatFH56**
ATGAGCCTGCCCTGGGTCTGCTAACCTCCACATGCAGCATGGATCTCCCAGGCAGCAGCAGC
CTGCTCTGGGTCTTTGCCCAGCAGC
**Primers**:
Forward: 5' GAGCCTGCCCTGGGTCTGCT 3`
Reverse: 5` GCTGCTGGGCAAAGACCCA 3`
**Oligoprobe**:
5` GCCCTGGGTCTGCTAACCTCCACATGCAGCATGGATC 3` biotin or digoxigenin

**SatFH213**
AGCCATCATATTCTCATTTACAACATTAAAGTAAATAAACAATTTAGACTTGAAATATTTTTC
TAATTTCTTAATAGGGCTCTGGAAGGGAAATTTGGAAACAGATTTAATAATAATTTTAACAT
TATATTAAAATAAGTGGCGTATTTTTTTTTTTAGGTCCTAAATGTACATAACGCGGTAGCGTA
CGTACATATAAAGTGTACATATCGCGGTAGGTTACGGGCTGACTCATTGGAAATCAAGCTCC
ACTGTTGTGTCTGAAACTGCACTACACGACCACAACTACCGGGGGAGGGGAACAGCTTTTGC
TATTGGCTGAGAGCGTGGCCTATGTAATTATGTGGGTTAAGGAGGGGTCAAGGTGTACAGA
AGGAGGGAGGCTGTACAGGCAACATGGAAGTACAAAACGGCTCTCTGCCCCCTATATATAG
GGGCTAAGGGGTGTAGCAGAGTGGATAACAACATTGCCTTCCACCCAGAGGAGCAGGGTTC
GAATCTCACCTC
**Primers**:
Forward: 5' ACGGGCTGACTCATTGGAAA 3`
Reverse: 5` TGAGATTCGAACCCTGCTCC 3`
**Oligoprobe**:
5` GTGTACATATCGCGGTAGGTTACGGGCTGACTCATTGG 3' biotin or digoxigenin

**SatFD150**
AACCAGAGCCAGGCTCAGTCCAATTCGCCTCCAAATCCATCGTCTTTCAGTACTGAACCACT
TCCTCAGACACCTCACATCTCCAGGGCAGGTGTAACCACCGGAGACACCGGGAAATGCATC
CCGCACACGTCTGGTAAGGCTCCTAACAGGCACATGCAACCAAAACACTTGGAGCAGTTTGC
CATCGTTTTGGGCAAGTTCCTGAAGCCAGACAAACGAGGTGTCTGCGGACTCAGGGGGAAC
CCCTGAGCATGCACGACCGACTCCAAGCTCCGTCGGACGCTTCCCGGCCCGCCCTCCGCCCC
AAACGCCCGCCGCTGAGGATGGGACATTTCTCAAACGTCGCCCCAGACGTGTCGTGTGGGGC
GTCTGGCGAAGCGGCACGAGCCCCCGAGCGCAACGGCGGGCCTCCTTTCCCCGTGCCGGGCT
CCCCTCTTCGATTTCGACCGTTTTGTCCGATCCTTTTTGCGCCGTTTCGGCCTACCGCCAGAA
CCGTCCGTCGCAGAGAGACGCCGACGCCTCAGGCGCGTCCGCAGCCCCCCCAAACATGGGG
GAGGAGCCCGCTTTCAGCTCCCTACCGCCTTTACTTTACCCCCAAATCCACTTTTGTCCGATC
CTCAGGTGGTGTTTACCCCCTAACCTAAGACGGATTAGTGCAACGCCTCTTTTGGCCAGCAG
ATGGAGCTGGCTCTCTCCTTTAAACAGTTCCCTGCTGTGGCCAGCAGGTGGAGCTGTATCCT
GCCCACTCAGTCTATGCAGTGGGAAAACATATGTTCCTTGGCAAGCAGAGCCTCACCCTAAC
AGTTCCCTGCTTTGGCCAGCAGGGGGGGAAATCTGCTGTTCCAGCTTCATGCACCCTATCAG
GCTGAAATTCAAAGCTTCTGGCCAGCAGAGAAGGCTCAAATTGGGCACACATCATCTTCAGG
GTGCCTTTGACGCATTCTGCCACTTTGAAAGGGCTCGCTGCTCTACCTTTCGAGTGAGATTTT
TTGCAAAGTGTGCACCTTCCGCGGCCGCTAGCGGGCGCTACAGTGGGCCCACGATGCCGAA
ATTGCTCTGAAAAATAATTGAGTGTGTGCCCCACATGCCTGCCGAGTTTCACAGGCCTAGCT
GTCTCCGTGTTGAAGTTGTGAACGTTTTAACTTTGGGTCTCATTTGGCAAAATGGTGCAATGC
ACGTGTTGGCCAGCAGGTGGCTGGACACTCACGCTATGCTGTGCTCCCACACATATGTCTTT
CCATACACTCCACTATCACAGCCTCACCCCAACGCTTGCCAACAGCAGAGATTCAACTCTGT
GCACTCAGACCAGGCCGCAGAGCGACTCGTCTGCTCTTTGGGAAGCGTGGGGAATAAAAGC
AGGCCTCA

**Primers**:
Forward: 5' TTGCCATCGTTTTGGGCAAG 3`
Reverse: 5` GGGAACTGTTAGGGTGAGGC 3`

**Oligoprobe**:
5` GGAGCAGTTTGCCATCGTTTTGGGCAAGTTCC 3' biotin or digoxigenin

**SatFD233**
TGAAAAAACCCGGCTATCCATTATCTCATGATAACCATGTCATTAACTCAAGAAAACTGTTA
TTGTTTTCTCGTGATAACAACATAGCTAAGTCATTATCTGGAGAAAACAATGTAAAAGAAAG
CATTCAAATACACCCGCTCTCAGCTTCCGTAGCTCATCACTGGTTCCAGGAAAATGAAAAAA
CCTGTTGCCTTTAAAGCATTTCTTTCTTTCCTTTTACATTTGAAGGACTTGTATTTCTCATCAC
TACAAATTGTGTCTATCTATGCCTAAACTGAGAGCCACTGAGTTTATCTCACACATACTTGTT
GTTAGAGCAGTTCAGGTCTTAAATGTCCCAACACAGAAAGCGTTTTAGTTATAGAAAGGTTA
TTACTGACCACCTTTATGATGGATTCAACTAATCTGCTTTCAAAGACACCGTTTAACTAAATT
AAGCGACAAACACTCAGAAAGTTCAACGACTGTTGATTAAGTTAGTAAAAAATGTTTTAATA
AAAATAGTTATTACCAGCAATTAAATAATAGTTGTACTTCAGCACTATTATAGAAAATAACG
ACTCCATCAAGGCCCAGTCCAAAACAACATCACCATTTTCTGTAAATCTAGATTTCTGTTCAC
TTCTACGTGACTTTACCACGTAGCACATTCACTCTAACTCCAGAAGTGTTTCTTTAATAGTTT
TTTCTTTGTTTTTTTAGTTAAACAAACTTCAGTGAACTTCAAGTGTTAAGAACACAAAAACAC
AGAAACCATGACAGACTTGGAAAGAACTGAAATCTACCTTTTTTCCATCCTGAAAGTTTCTA
GTTTGACTTTTGTCAGTTTCAGAGACAATAATTGTTGTCAGGAAGCTGGACATCTTTAAGAG
AAGCGATTTTTATGGCAAAATAATCAAATAAGGTCTGAGAAGGCTTTAAGGTTGGATCTGGC

TGGTGTTTAAACGATGCTGCCATGATGCACCTAAGTGGGAACCATAGCAACCATGGCCTTCA
CAGCATATTTCTTAAAATCAGCTGTTTTAAAACATAAAAAAATAAAAAGCCAGATCTAGGTT
TCTACACAAACCTCATTACGGAAGCCGAGAGCGGACCGGTCCCTCCCATACTTTTTGATGAT
GGTTTTCTCGTTATAACGAGTTAATTTTCTCCTTTTCTAAAGACAACAAAGGCCGTTTTCTTG
TGATAATGGGATGATTTATCTGGAGAAAACAAACTTTGTCGGGCAGTCGTCTCCTTAAATGA
CGAGGCTGATATAAAAAGTTATCTAGAAGACAGACAAATTGCAATTTCAGCCTGTGTTAGAC
CTGGAGAGAAGTAAAATCTGATTCGTTGATCATGATCATGTCAGATCGTGGTTCTCCACCAT
CTGACATTTTCAGCGTGAGCCTTTTCTGGTAAGGCTTTAAGCACACTGAGAAACTGAACATA
AGCATTAAAAAAACCC

**Primers**:
Forward: 5'CAAATACACCCGCTCTCAGC 3'
Reverse: 5`TGTTTTGGACTGGGCCTTGA 3'

**Oligoprobe**:
5` CAGAGACAATAATTGTTGTCAGGAAGCTGGACATC 3` biotin or digoxigenin

**SatFD95**

GCCCAGCCAGAGGCCCTCTTCTGGGAAGCTCTGTGACACAGCCCCCTTGCTGGCTGGAGGTT
TTTTTTTTTCTTTCTTTCTTAATCCAGGGGAAATTCTTTATTTAATTTTGCTATCACCCCCATGC
CAGGAACTTTGAAGAAATTTCATCCAGGGCAGGGGACCTTTGCTGCACTACTG

**Oligoprobe**:
5` CATCCAGGGCAGGGGACCTTTGCTGCACTACTG3` biotin or digoxigenin

**SatFD207**

CTCAACTTGTTACATCAGATGGTCACGAGTTCTTTACGTTATGACATCAGATAGCCATTCTGT
TCTCACCTTATGACATCAGATTGTCATGCTGTTCTTCGCCTTATGACATCAGATAGTCATTCC
GTT

**Oligoprobe**:
5`CATCAGATGGTCACGAGTTCTTTACGTTATGACATC 3` biotin or digoxigenin
